# A *DNM1L* GTPase Effector Domain Deletion Causes Intermediate Defects in Peroxisomal and Mitochondrial Dynamics and Reveals a Targetable Metabolic Vulnerability

**DOI:** 10.64898/2026.11.03.763232

**Authors:** Saurabh Srivastav, Sharayu V. Jangam, Jonathan C Andrews, Evangelos Theodorou, Lauren C. Briere, Gabriel D Quintana, Ingrid LS Gomes, Jonas A Saute, Pranjali Bhagwat, Aanya Subramaniam, Neelanjana Roy, Ellen Macnamara, Amarilis Sanchez-Valle, Cecilia Gordillo, Undiagnosed Diseases Network, Hugo J Bellen, Shinya Yamamoto, Seema Lalani, Michelle Duffe, Frances A. High, Melissa A. Walker, Leonardo Simão Medeiros, David A. Sweetser, Michael F Wangler

## Abstract

DNM1L encodes dynamin-related protein 1 (DRP1), a key GTPase regulating mitochondrial and peroxisomal dynamics. Pathogenic *DNM1L* variants are associated with a broad spectrum of neurodevelopmental and neurodegenerative disorders, with phenotypic severity varying by affected protein domain. While variants in the GTPase domain are typically associated with milder phenotypes and middle-domain variants with severe developmental and epileptic encephalopathy, the functional impact of GTPase effector domain (GED) variants remains less defined. Here, we describe four individuals harboring a recurrent heterozygous GED deletion (L700del) presenting with movement disorders and lacking severe infantile encephalopathy, consistent with an intermediate clinical phenotype. Patient-derived fibroblasts exhibited tubular peroxisomes and enlarged mitochondria, while *Drosophila* models demonstrated that *DNM1L^L700del^* has intermediate functional effects that were more pronounced than those of a GTPase-domain variant A192E but less severe than those of the middle-domain variant G350R. Comparison with another GED variant, Y691C, further demonstrated variant-specific differences in severity within the GED. Metabolic stress assays showed that L700del fibroblasts exhibited impaired growth under galactose conditions, consistent with increased vulnerability under conditions dependent on mitochondrial oxidative metabolism. Bezafibrate, a PPAR agonist that promotes mitochondrial biogenesis, partially rescued this growth defect. Together, these findings support a functional allelic spectrum in *DNM1L*-associated disease in which protein domain contributes to, but does not completely determine, functional severity. L700del represents an intermediate-strength allele associated with a movement-disorder phenotype and a pharmacologically modifiable metabolic vulnerability.

## Introduction

The *DNM1L* gene encodes Dynamin-related protein 1 (Drp1), a large GTPase that is critical for the fission of both mitochondria and peroxisomes(1, 2). Drp1 orchestrates membrane scission of mitochondria and peroxisomes by assembling into oligomeric spirals at sites of fission. The process of fission itself is mediated by GTP hydrolysis (3, 4). While Drp1’s role in mitochondrial dynamics has been extensively characterized, its involvement in peroxisomal division is increasingly gaining attention, underscoring its dual-organelle regulatory function (5–7).

Structurally, Drp1 contains three conserved regions the GTPase domain (GD), middle domain (MD), and GTPase effector domain (GED) which contribute to GTPase activity, oligomerization, and regulation of membrane fission (8). Disease-associated variants in these domains produce distinct functional and clinical consequences. For example, pathogenic GTPase domain variants can impair GTP binding or hydrolysis and some have been linked to autosomal dominant Optic Atrophy (OPA5) (9). In contrast, heterozygous MD variants, frequently exert dominant-negative effects, and are associated with severe neurodevelopmental disease characterized by mitochondrial elongation, defective peroxisomal division, refractory epilepsy, and early lethality (5, 7, 10). GED variants are considerably less characterized having been reported in individuals with neurological phenotypes including developmental delay, epilepsy, hypotonia, and encephalopathy(7). While all the variants discussed above are typically heterozygous *de novo*, there are recessive forms of *DNM1L* related disorders that have been reported (11, 12).

These emerging genotype-phenotype associations suggest that the location and functional severity of *DNM1L* variants may influence both organelle dysfunction and clinical phenotype (10, 13, 14). However, the relationship between affected protein domain, degree of functional impairment, and disease severity remains incompletely understood, particularly for GED variants. *Drosophila melanogaster* provides a powerful system for addressing this question *in vivo*. Loss of *Drp1* in *Drosophila* disrupts mitochondrial fission and distribution, impairs synaptic function, and causes lethality (15). The loss-of-function alleles *Drp1¹* and *Drp1²*, originally identified through forward genetic screens, produce profound mitochondrial and synaptic defects and provide a sensitized genetic system for assessing DRP1 function *in vivo* (15). We have previously used Drosophila to model disease-associated *DNM1L* variants (7, 16). Importantly, variants associated with distinct human phenotypes can be directly compared within a common genetic background. These include A192E, a GD variant associated with autosomal dominant optic atrophy; G350R, an MD variant associated with severe encephalopathy and epilepsy; and Y691C, a GED variant associated with neurological disease and abnormal mitochondrial and peroxisomal morphology. This approach allows us to test whether the striking clinical heterogeneity of *DNM1L*-associated disease reflects a corresponding spectrum of functional effects on mitochondrial and peroxisomal dynamics.

Here, we describe four individuals with a recurrent *DNM1L* GED deletion, L700del, associated with an early-onset movement disorder characterized by dystonia, tremor, and gait abnormalities without severe infantile epileptic encephalopathy. Using patient-derived fibroblasts and *Drosophila* models, we show that L700del disrupts mitochondrial and peroxisomal dynamics and exhibits an intermediate functional severity compared with disease-associated GD and MD variants. L700del fibroblasts also demonstrate a selective growth defect under metabolic conditions that increase dependence on mitochondrial oxidative metabolism. Based on this bioenergetic vulnerability, we tested bezafibrate, a PPAR agonist that promotes mitochondrial biogenesis and oxidative metabolism, and found that it partially rescued the cellular growth defect. Together, these findings support a functional allelic spectrum of *DNM1L*-associated disease and identify a metabolic vulnerability that can be pharmacologically modulated.

## Material & Methods

### Human Subjects

All four research participants provided informed consent for study participation and publication of clinical information in accordance with protocols approved by the relevant institutional review boards or ethics committees. Subject 1 was evaluated and enrolled in the Undiagnosed Diseases Network at the Harvard clinical site. Subject 2 was evaluated at Baylor College of Medicine and was enrolled in an IRB approved protocol focused on neurodevelopmental disorders (H-44779). Subject 3 had been enrolled in the Undiagnosed Diseases Program (UDP) study prior to its expansion into the UDN. Subject 4 was identified at a Brazilian referral center for rare diseases after detection of the same *DNM1L* variant and subsequent contact with the UDN through ClinVar. Consent was obtained according to local institutional procedures and included permission for publication of clinical information and photographs.

### Cell Culture

Human skin fibroblasts from subjects harboring the *DNM1L^L700del^* heterozygous variant and reference control were cultured in Dulbecco’s Modified Eagle Medium (DMEM) supplemented with 10% fetal bovine serum (FBS) and 1% penicillin-streptomycin at 37°C in a humidified incubator with 5% CO₂.

### Immunostaining

Cells were seeded on coverslips, fixed with 4% paraformaldehyde, and permeabilized with 0.5% Triton X-100. The cells were separately stained for peroxisomes and mitochondria using primary antibody against human PEX14 (Abcam ab183885, 1:500) for immunostaining peroxisomes and with anti-ATP5A (mitochondrial ATP synthase subunit alpha) (Abcam ab14748, 1:1000) for immunostaining mitochondria. The cells were washed three times (5 min each) with PBST followed by incubation with primary specific secondary antibodies. For Pex14, donkey anti-rabbit IgG conjugated to Cy3, (Jackson ImmunoResearch Laboratory 711-165-152), at a dilution of 1:1000 was used. For anti-ATP5A, anti-mouse cy3 conjugated secondary antibody (1:1000) (Jackson ImmunoResearch Laboratory 711-165-151) was used. Further, Phalloidin -488 (A12379) and DAPI were used to visualize actin cytoskeleton and nuclei, respectively.

### Fibroblast Cell Imaging

Cell samples were mounted using Vectashield antifade reagent (Vector Laboratories: H-1200-10). The samples were imaged using a Zeiss LSM 800 confocal microscope equipped with Airyscan. The 63x water objective was used for image acquisition, and identical settings were used for all groups to ensure imaging consistency.

### Fibroblast growth assay under metabolic stress

Fibroblasts from individuals harboring the L700del variant and control fibroblasts were maintained in DMEM supplemented with 10% FBS, 1% penicillin-streptomycin, 1% non-essential amino acids (NEAA), 1% GlutaMAX, and 5 ng/mL basic fibroblast growth factor (bFGF). For metabolic stress experiments, 130,000 cells per well were seeded in 12-well plates in triplicate and cultured in glucose-free DMEM supplemented with 10% FBS, 1% penicillin-streptomycin, 1% NEAA, 1% GlutaMAX, and 5 ng/mL bFGF, with either 10 mM glucose or 10 mM galactose as the carbohydrate source. Mutant and control fibroblasts were cultured in the presence or absence of 100 μM bezafibrate. Six days after seeding, viable cells were quantified by Trypan Blue exclusion.

### *Drosophila DNM1L* transgenic stock generation

We had previously generated transgenic flies with human *DNM1L* Reference and a p.G350R variant (16) (*DNM1L^Ref^* and *DNM1L^G350R^*, respectively) and human *DNM1L* p.Y691C variant and a GED domain deletion construct (7) (*DNM1L^Y691C^* and *DNM1L^ΔGED^*, respectively). For this study, we produced GD domain variant *DNM1L* p.A192E (*DNM1L^A192E^)*, and a GED domain variant *DNM1L* p.*L700del* (*DNM1L^L700del^*). Human cDNAs used for the generation of *Drosophila* transgenics were codon-optimized for *Drosophila* expression (GeneArt, ThermoFisher). Constructs were subcloned into the pUASt-attB vector. Transgenic flies were produced by injecting prepared DNA into VK00033 embryos, and the site for site-specific integration was (*y[1] w[1118]; PBac[y[+]-attP-3B]VK00033*) (17).

### *Drosophila* Genetics and Peroxisomal Morphology Analysis

The *Drp1^1^* and *Drp1^2^* loss of function (LOF) alleles used in this study were originally described by Verstreken et al. (2005) (15). Flies carrying the *Drp1^1^* and *Drp1^2^* alleles and human *DNM1L* constructs were generated through a series of crosses within our lab.

To examine peroxisomal morphology, a peroxisomal reporter was expressed in the third instar larval salivary gland. A UAS-GFP-SKL construct encoding a GFP tagged with a carboxy-terminal SKL (peroxisomal targeting signal 1) was cloned into the pUASt-attB vector. The transgenic insertion on the second chromosome was recombined with the *Actin-GAL4* driver line (*y¹ w⁎; P{Act5C-GAL4}25FO1/CyO, y⁺*) as described by (18), enabling robust, ubiquitous expression of GFP-SKL.

Transgenic flies with *UAS-DNM1L^Ref^, UAS-DNM1L^A192E^, UAS-DNM1L^G350R^, UAS-DNM1L^Y691C^,* and *UAS-DNM1L^L700del^*were expressed in *Drp1^1^* mutant background using *Mef2 Gal4, daughterless-Gal4 (da-Gal4), elav-Gal4, Repo-Gal4* driver transgenic lines.

### Drosophila Tissue Immunohistochemistry

Wandering third-instar *Drosophila* larvae were collected for immunohistochemical analysis of the larval brain and body wall muscle. Larvae were dissected under a stereomicroscope on Sylgard-coated dishes in 1× PBST (phosphate-buffered saline [PBS] containing 0.1% Tween-20). For body wall muscle analysis, ventral longitudinal muscles 6 and 7 in abdominal segment A3 were examined.

Following dissection, tissues were fixed in 4% paraformaldehyde (PFA) in 1× PBS for 10 minutes at room temperature and washed three times in PBST for 10 minutes each. Samples were blocked in 5% normal goat serum (NGS) in PBST for 30 minutes at room temperature. Tissues were incubated overnight at 4°C with primary antibodies diluted in blocking buffer, washed three times for 10 minutes each in PBST, and incubated overnight at 4°C with the appropriate fluorescently conjugated secondary antibodies. Samples were then washed three times in PBST and mounted in VECTASHIELD antifade mounting medium containing DAPI.

Peroxisomes were labeled using rabbit anti-Pex3 (1:500; McNew laboratory, Rice University) (19), followed by Cy3-conjugated donkey anti-rabbit IgG (Jackson ImmunoResearch, 711-165-152; 1:1000). Mitochondria were labeled using mouse anti-ATP synthase subunit α (Abcam, ab14748; 1:1000), followed by Cy3-conjugated donkey anti-mouse IgG (Jackson ImmunoResearch, 715-165-151; 1:1000).

Fluorescence imaging was performed using a Zeiss LSM 800 confocal microscope equipped with Airyscan and high-numerical-aperture water-immersion objectives. Z-stacks were acquired from larval brains and ventral longitudinal muscles 6 and 7 in abdominal segment A3. Airyscan reconstruction was performed using ZEN 2.6 software.

Image analysis and quantification were performed using Imaris software (version 9.8.2; Bitplane, Zurich, Switzerland). Puncta quantification and morphological analyses were performed using the Surface module as described in the Image Analysis section.

### Fluorescence microscopy imaging

High-resolution imaging was performed using a Zeiss LSM800 Airyscan confocal microscope (Carl Zeiss Microscopy GmbH, Germany) equipped with Plan-Apochromat 63×/1.2 NA or 40×/1.2 NA water immersion objectives. Images were acquired with a frame size of 1024 pixels to ensure optimal resolution. Fluorophores were excited and detected using the following laser lines and spectral ranges: 405 nm excitation with 410–480 nm emission for DAPI, 488 nm excitation with 493–570 nm emission for Alexa Fluor 488, and 561 nm excitation with 576–700 nm emission for red fluorophores such as Alexa Fluor 568 or 594.

Raw image data were processed using the Airyscan Processing module in ZEN 2.6 software (Blue edition). The 2D Super-Resolution (SR) mode was applied to enhance spatial resolution beyond the diffraction limit. Deconvolution was carried out using the built-in Wiener filter, with the filtering parameter set to ‘Standard’ to balance signal restoration and noise suppression. All processing settings were maintained constantly across all experimental conditions to ensure consistency and comparability of results.

### Image quantitative analysis

Quantitative analysis of fluorescence images from cells immunostained with anti-Pex14 and tissue sections stained with anti-Pex3 was performed using the Surface module of Imaris software (version 9.8.2; Bitplane, Zurich, Switzerland). In this module, 3D surfaces were rendered for the specific fluorophore signals based on intensity, using the background subtraction option to enhance signal specificity. To isolate individual puncta corresponding to peroxisomal structures, an area-based filtering strategy was applied. A lower threshold limit of 0.5 µm² was set, whereby only surfaces with an area equal to or greater than 0.5 µm² were retained for further analysis.

To exclude nuclear-localized signals from the quantification, particularly important in the case of Pex3, the surface masking tool within the Imaris Surface module was employed. First, a separate surface was created for the nucleus using the DAPI channel. This nuclear surface was then used to mask the Pex3 channel by setting all voxels within the nuclear volume to zero, thereby eliminating any contribution from nuclear-localized Pex3 signal in the final quantification.

The total number of surfaces generated post-thresholding corresponded to individual peroxisomal puncta and was extracted from the statistical output generated by Imaris.

Mitochondrial morphology was analyzed using ImageJ (NIH). The width of individual mitochondria was measured and quantified in both patient-derived cells and Drosophila tissue samples, enabling comparative evaluation across model systems.

### Larval Tracking

Larval locomotion was assessed in wandering third-instar larvae according to a previously established protocol (20). Briefly, 1–4 larvae were placed on 100 × 20 mm polystyrene plates containing 250 mL of solidified grape juice medium (1.8% agar, 45% grape juice, 2.5% acetic acid, and 2.5% ethanol). Larval movement was recorded using a Basler 1920UM 1.9-megapixel monochrome USB3 camera and Basler pylon software at an acquisition rate of 33 frames per second.

Videos were analyzed in ImageJ using the wrMTrck plugin with the optimized third-instar larval detection parameters described previously (20). A Kalman Stack Filter was applied prior to tracking, and larval velocity was quantified using wrMTrck. Videos in which larvae contacted one another were excluded from analysis to avoid tracking errors or incorrect assignment of individual trajectories. Quantitative data were analyzed using GraphPad Prism.

### Pupal Length

Pupae were collected and imaged with a ruler included in each image for spatial calibration. Images were analyzed using Fiji/ImageJ (NIH), and the image scale was calibrated using the ruler. Pupal length was measured along the central anterior-posterior axis of each pupa. Measurements were analyzed using GraphPad Prism version 10.

### Statistical Analysis

Statistical analyses were performed using GraphPad Prism version 10. Comparisons between two groups were performed using two-tailed unpaired Student’s *t* tests. Comparisons involving multiple experimental groups were performed using one-way ANOVA followed by Dunnett’s multiple-comparisons test. Data are presented as mean ± [SEM], unless otherwise indicated in the figure legends. The number of biological replicates (*n*) and specific statistical tests used for each experiment are provided in the corresponding figure legends. A *P* value < 0.05 was considered statistically significant.

## Results

### Clinical Cases

Through the Undiagnosed Diseases Network (UDN) and GeneMatcher, we identified four individuals with previously undiagnosed neurological disease characterized by early-onset movement disorders (**Table 1**). All four subjects presented in infancy or early childhood with difficulty learning to walk, wide based-gait and gross motor delay. All four subjects also exhibit dystonia, and tremor and three individuals showed a partial response to L-dopa. Additionally, two subjects exhibit axonal sensory neuropathy. Of note, three subjects had no seizures, and two had normal brain MRI. Clinical details on these subjects reveal significant overlap (**Table 1**).

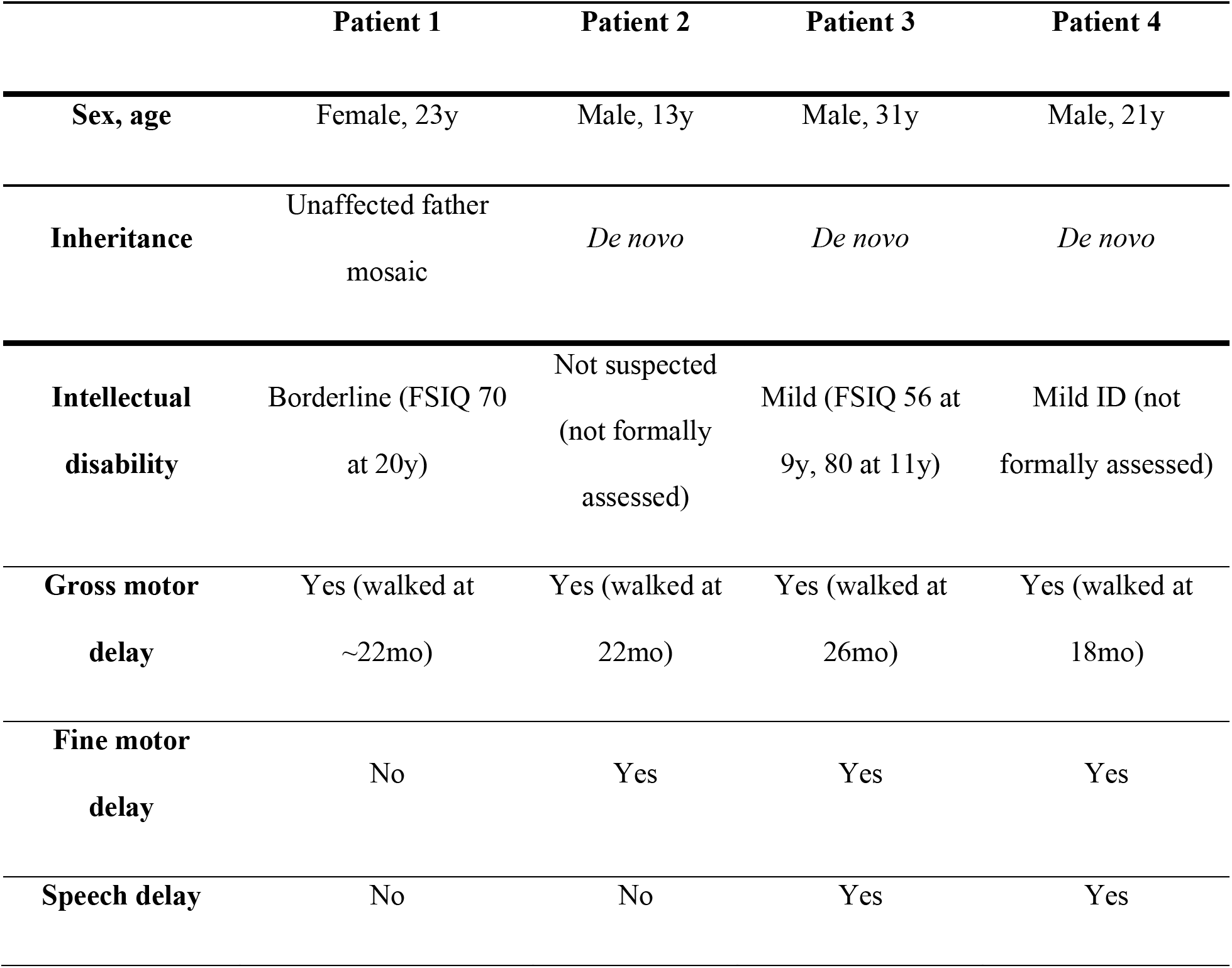
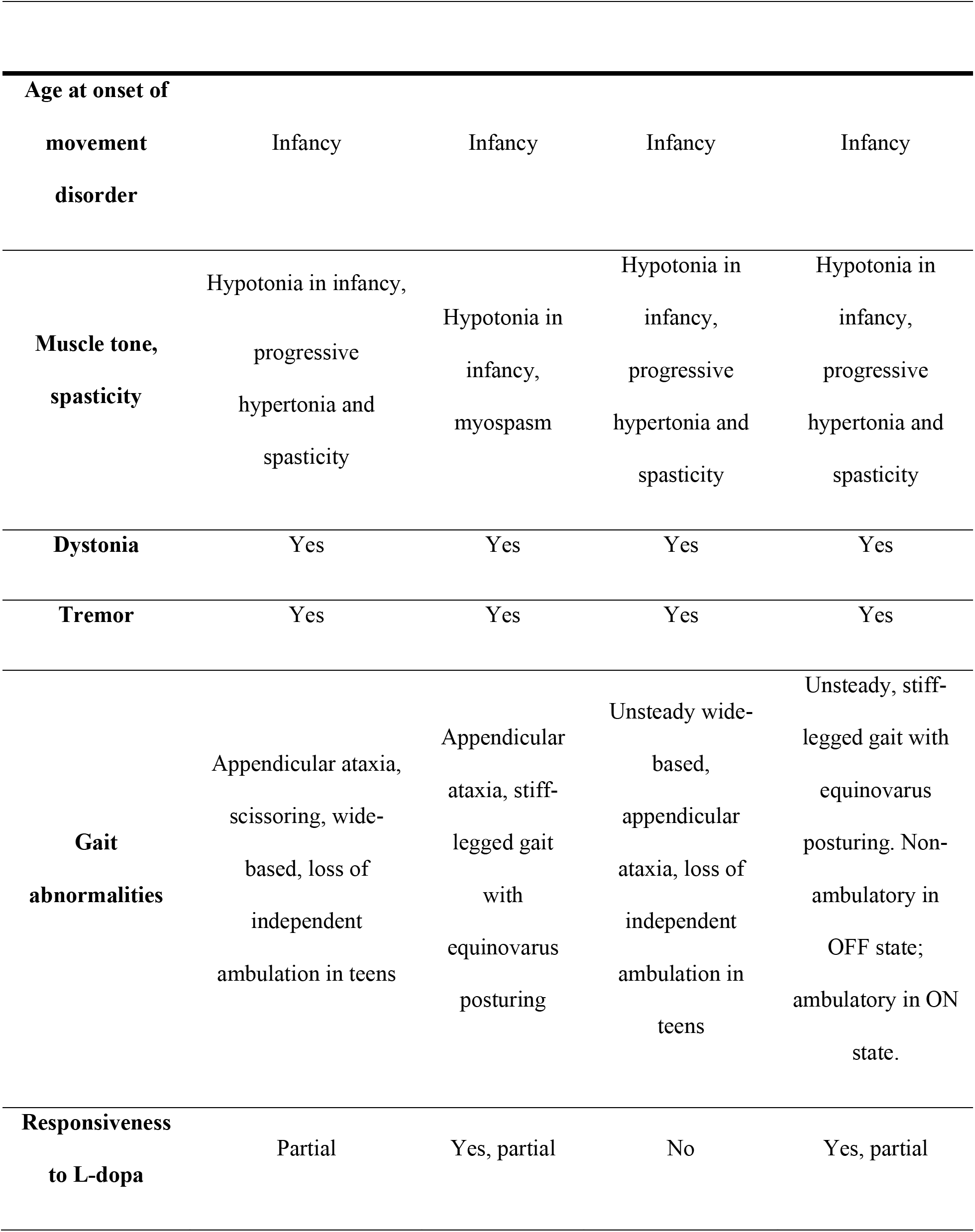
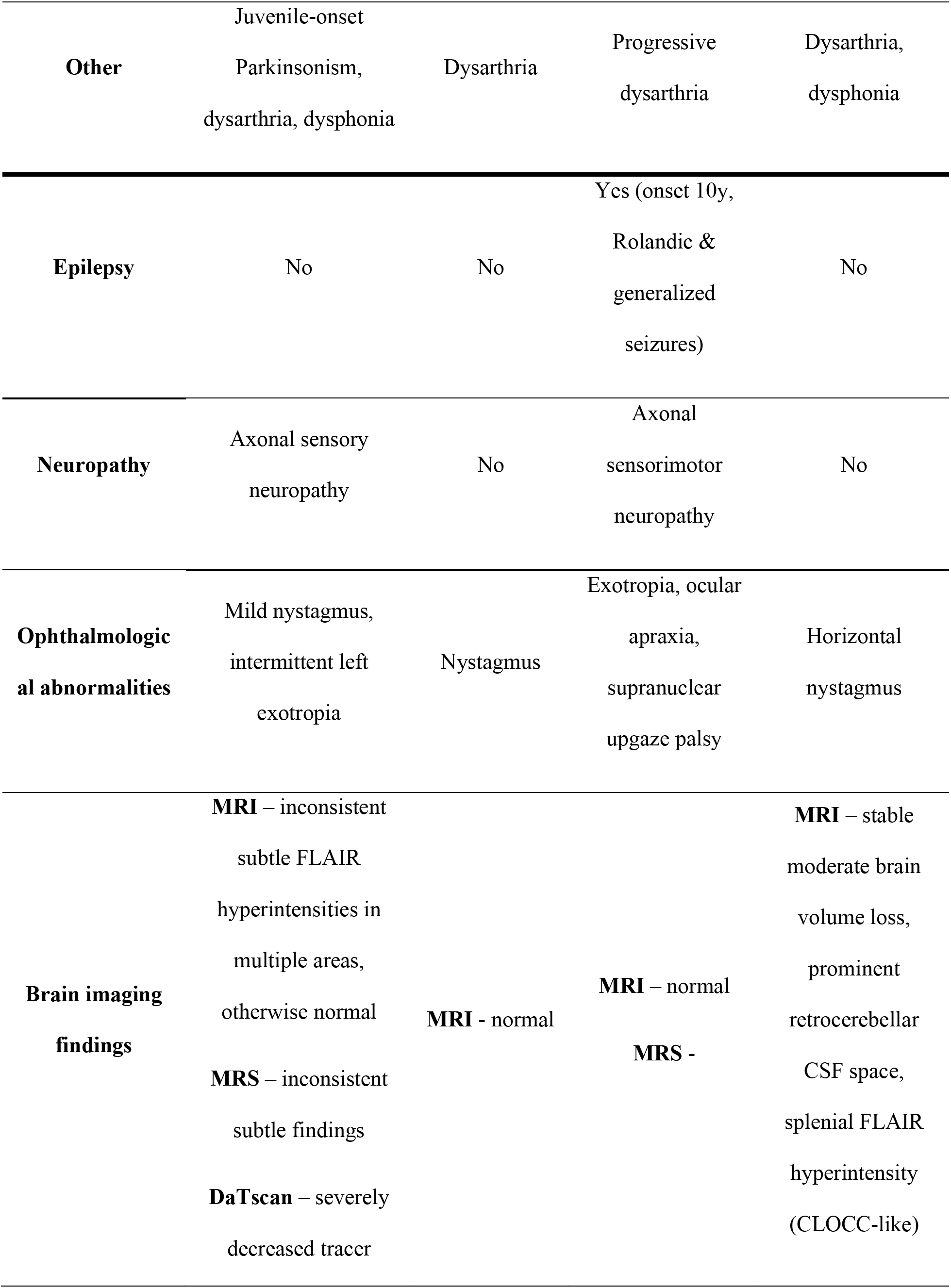
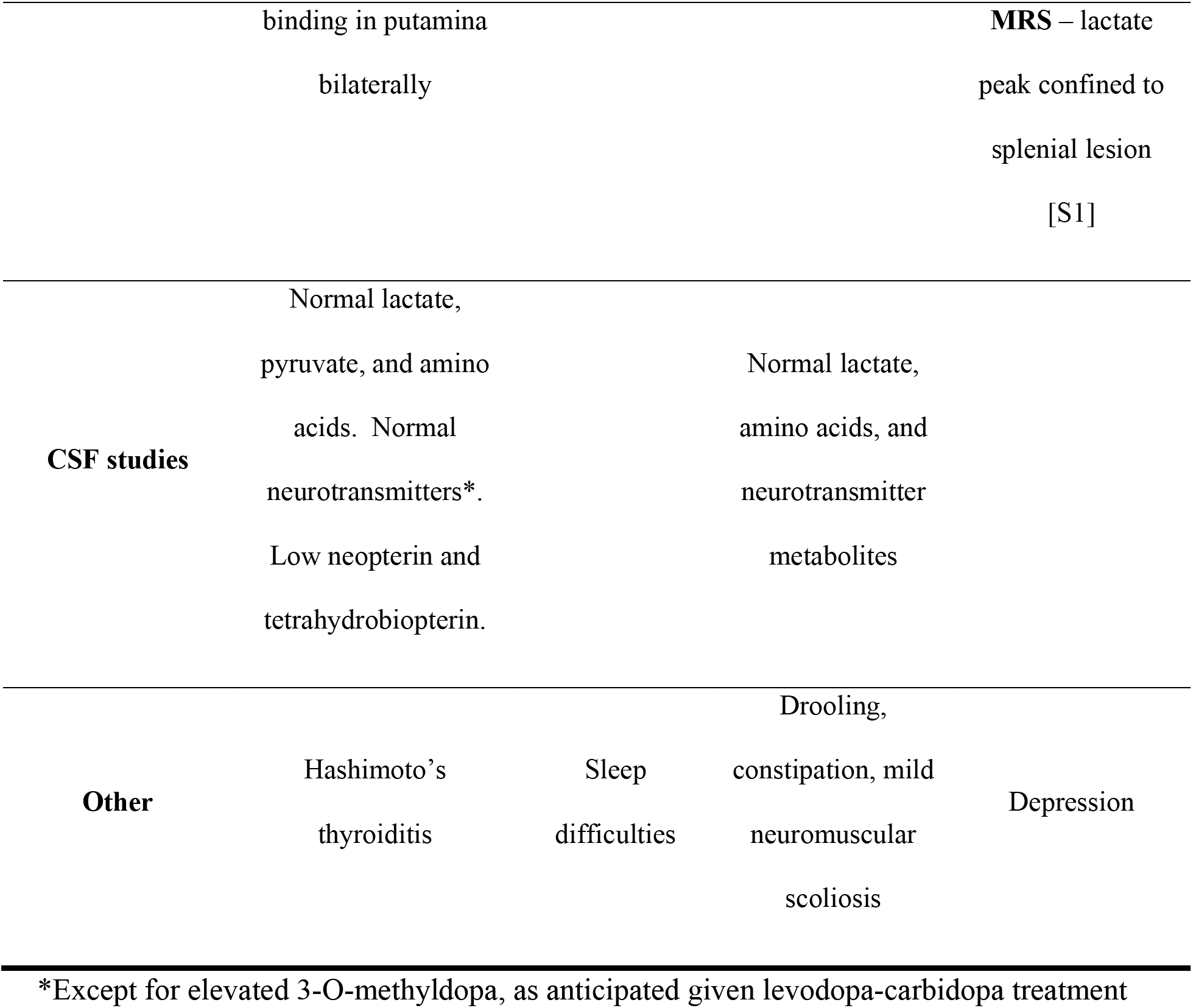

All four subjects share the same heterozygous in-frame deletion in the GED of *DNM1L* (**Fig 1A & 1B**) (NM_012062.3: c.2097_2099delTCT, p.(*L700del*)). The variant arose *de novo* in Subjects 2-4 and the father of Subject 1 had low-level mosaicism. All four subjects had substantial overlap in phenotype, but their phenotypes differ from those classically associated with DNM1L (OMIM #614388, Encephalopathy due to defective mitochondrial and peroxisomal fission, EMPF1). None of the four subjects exhibited infantile encephalopathy or experienced seizures during infancy; instead, their clinical presentation was dominated by early-onset dystonia, tremor, and gait abnormalities. Notably, L700del affects the GED rather than the middle domain in which many severe dominant *DNM1L* variants have been reported. The recurrent L700del variant and shared movement-disorder phenotype therefore suggested an expansion of the *DNM1L*-associated phenotypic spectrum and prompted functional investigation of its effects on DRP1 function.

**Fig 1:**
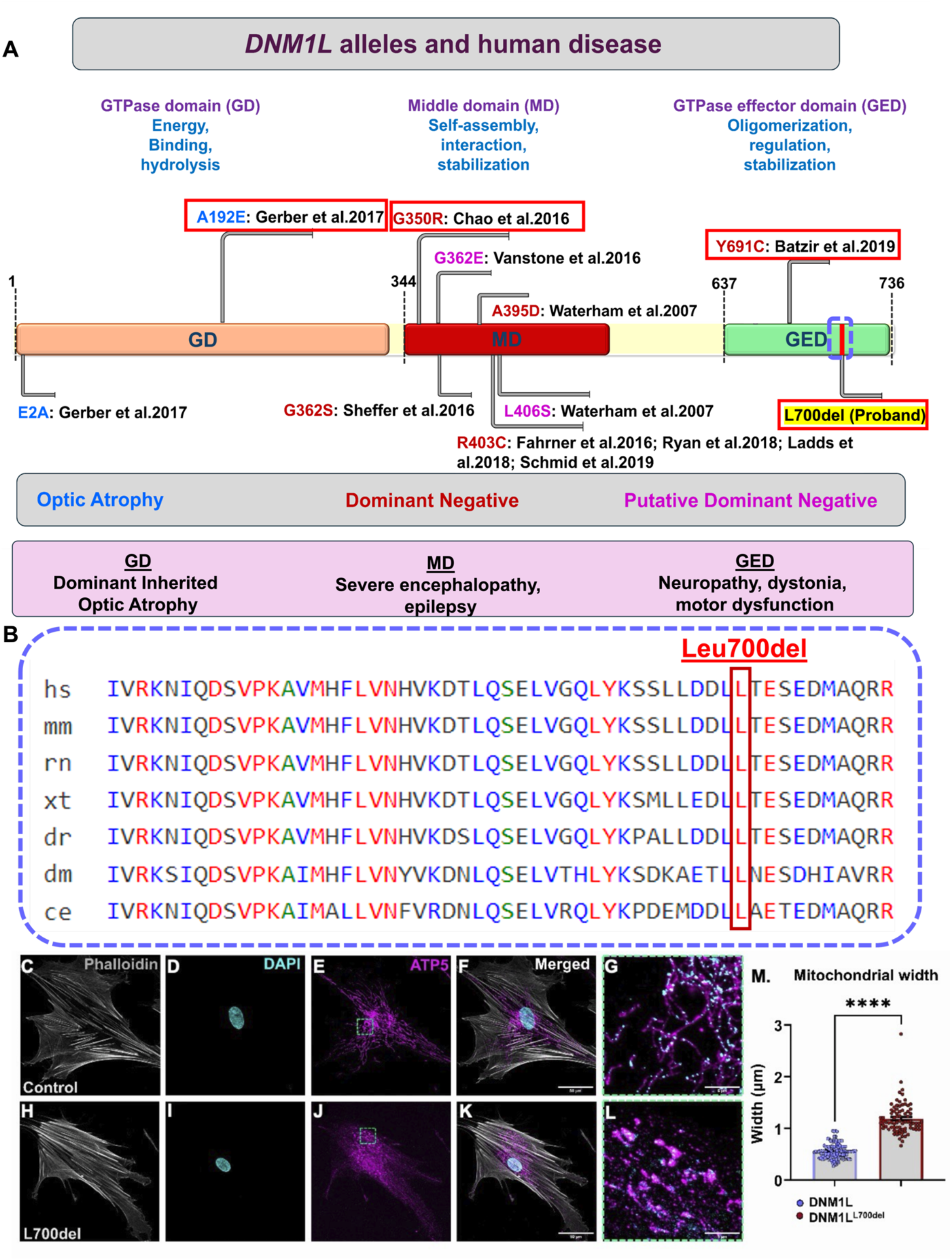
**A.** Schematic illustration of *DNM1L* genes and the protein functional domains linked to different disease-causing variants. **B.** Alignment of the L700del variant in human and model organism. **C-M.** Assessment of mitochondrial morphology in *DNM1L^L700del^* variant. Mitochondrial morphology was analyzed via ATP5 immunostaining in fibroblast obtained from control versus patient with *DNM1L^L700del^*variant (C-G & H-L respectively). The mitochondrial morphology in control appeared well distributed and connected (E & G). However, the mitochondrial morphology in *DNM1L^L700del^* variant appears swollen bulb like at several places with fragmented distribution (J & L). The mtDNA distribution was uniform throughout the mitochondrial network in control (G) unlike *DNM1L^L700del^* variant where it was mostly accumulated in those swollen bulb like structure (L). The mitochondria width was quantified using FIJI, and *DNM1L^L700del^* variant exhibited significantly higher mitochondrial width compared to control (M). Student t-test was performed for statistical significance ****p<0.0001

### The *DNM1L^L700del^* variant alters peroxisomal and mitochondrial morphology in patient fibroblasts

Because DRP1 regulates the fission of both peroxisomes and mitochondria, we examined the morphology of these organelles in skin fibroblasts derived from Subject 1.

To assess peroxisomal morphology, control and Subject 1 fibroblast cells were stained with phalloidin, DAPI, and an antibody against PEX14 (**Fig. S2**). Control fibroblasts contained predominantly spherical peroxisomes (**Fig. S2 C & E**). In contrast, Subject 1 fibroblasts exhibited a mixed population of elongated, tubular peroxisomes and spherical peroxisomes (**Fig. S2 C’ & E’**). Subject 1 fibroblasts also showed a reduction in total peroxisome number (**Fig. S2 F**), with a shift from predominantly spherical peroxisomes in control cells toward tubular peroxisomes in Subject 1 cells (**Fig. S2 G–H**). Thus, the L700del variant is associated with marked alterations in peroxisomal morphology and abundance.

We next examined mitochondrial morphology in control and Subject 1 fibroblast cells using ATP5A immunostaining (**Fig. 1C–1L**). Control fibroblasts displayed a tubular mitochondrial network distributed throughout the cell (**Fig. 1E, 1G**), whereas Subject 1 fibroblasts showed enlarged mitochondria with prominent perinuclear accumulation (**Fig. 1J, 1L**). Mitochondrial width was significantly increased in Subject 1 fibroblasts compared with controls (**Fig. 1M**), providing quantitative evidence of altered mitochondrial morphology. Subject 1 fibroblasts also exhibited an irregular distribution of mtDNA within the mitochondrial network (**Fig. 1Ls**).

Together, these findings demonstrate that the *DNM1L* L700del variant is associated with abnormalities of both peroxisomal and mitochondrial morphology in patient-derived fibroblasts.

### The L700del fibroblasts exhibit impaired growth under conditions requiring mitochondrial oxidative metabolism

To investigate the effect of L700del on mitochondrial metabolism, we compared the growth of patient-derived and control fibroblasts in medium containing either glucose or galactose, the latter increasing cellular dependence on mitochondrial oxidative metabolism. After six days in galactose-containing medium, fibroblasts from Subjects 1 and 3 exhibited significantly reduced growth compared with control fibroblasts, with reductions of 44% and 35%, respectively (**Fig. 2**). These findings demonstrate a selective growth defect under metabolic conditions that increase dependence on mitochondrial oxidative metabolism.

**Fig. 2:**
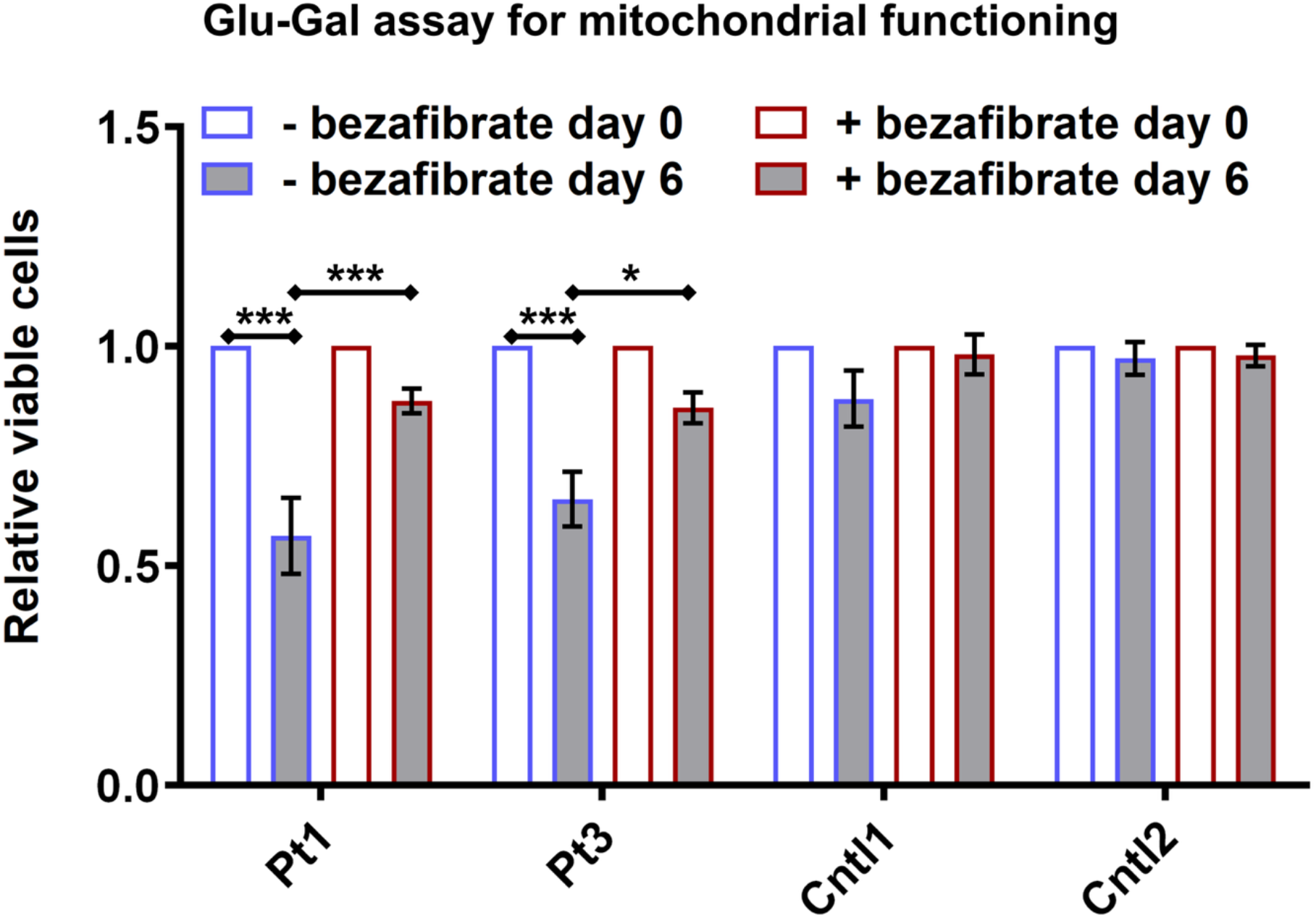
The growth of fibroblasts from patients 1 and 3 is impaired in galactose media relative to controls and is rescued by benzafibrate. Relative viable cell numbers of fibroblasts from two *DNM1L* patients [Patient 1 (Pt1) and Patient 3 (Pt2)] and two respective controls (Control 1 and Control 2) were measured after 6 days in 10 mM galactose medium, with or without 100 µM bezafibrate treatment at indicated time points (Day 0 and Day 6). Data are normalized to proliferation in 10 mM glucose. *DNM1L* mutants demonstrate reduced proliferation compared to controls when grown in galactose media. Bezafibrate treatment significantly improved viability in *DNM1L* mutant fibroblasts but had no effect on controls.

Next we tested whether this metabolic vulnerability could be pharmacologically modified. We administered Bezafibrate, an agonist of Peroxisome Proliferator-Activated Receptors that enhances mitochondrial biogenesis and oxidative metabolism via induction of PGC-1α. Treatment with this drug has been shown to improve cell viability of patient fibroblast with a middle domain variant (21). We therefore supplemented the cells with the Bezafibrate (100 µM). Bezafibrate increased viable cell numbers by 55% in Subject 1 fibroblasts and 31% in Subject 3 fibroblasts compared with untreated cells, while having no significant effect on control fibroblasts (**Fig. 2**). Thus, bezafibrate partially rescued the growth defect of L700del fibroblasts under metabolic stress.

### *Drp1^1^/Drp1^2^* exhibit altered peroxisomal and mitochondrial morphology

To establish the consequences of *Drp1* loss of function on mitochondrial and peroxisomal morphology *in vivo*, we examined *Drp1¹/Drp1²* trans-heterozygous larvae. These loss-of-function alleles cause severe defects in mitochondrial fission and result in larval lethality (15), but their effects on peroxisomal morphology have not been characterized.

In control larval body-wall muscle, Pex3-positive peroxisomes appeared predominantly as spherical puncta, whereas *Drp1¹/Drp1²* larvae exhibited elongated, tubular peroxisomes (**Fig. 3A, A’**). Quantification demonstrated a significant reduction in Pex3-positive puncta in *Drp1¹/Drp1²* muscle compared with controls (**Fig. 3C**). Mitochondria in control muscle formed an interconnected filamentous network, whereas mitochondria in *Drp1¹/Drp1²* larvae appeared enlarged and swollen (**Fig. 3B, B’**), with a significant increase in mitochondrial width (**Fig. 3D**).

**Fig 3:**
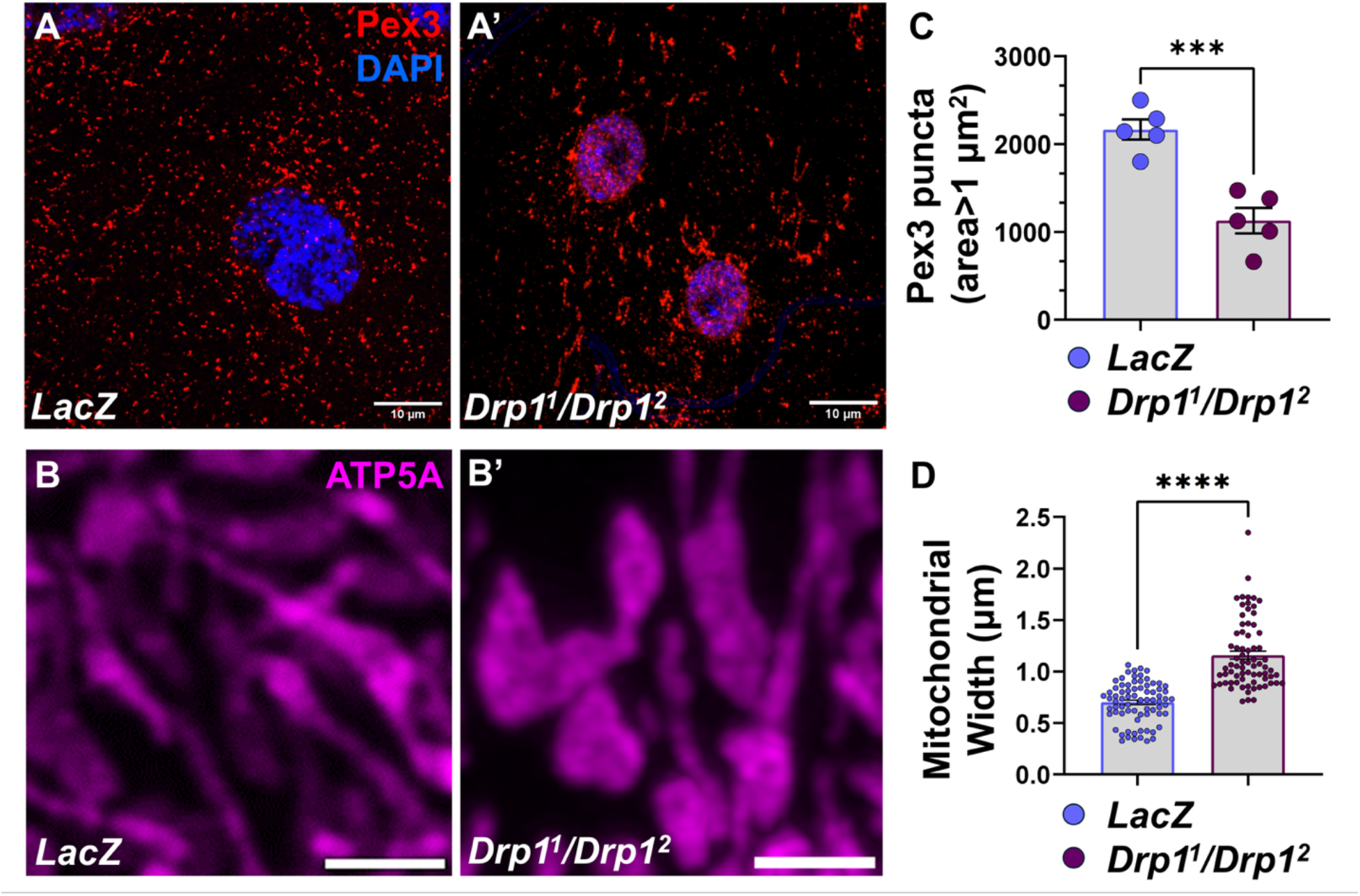
*Drp1^1^/Drp1^2^* exhibit altered peroxisomal and mitochondrial morphology. Panel A, A’ illustrates the pex3 stained peroxisomes from control and *Drp1^1^/Drp1^2^* respectively. Panel B, B’ depicts anti-ATP5a stained mitochondria in both control and *Drp1^1^/Drp1^2^*. The quantitative graph C illustrates peroxisomes number as pex3 puncta count in control versus *Drp1^1^/Drp1^2^* while graph D illustrates mitochondrial width comparison between control and *Drp1^1^/Drp1^2^*. ***p<0.001;****p<0.0001

Thus, severe loss of *Drp1* function disrupts both peroxisomal and mitochondrial morphology *in vivo*, producing organelle abnormalities similar to those observed in L700del patient fibroblasts.

### Expression of domain specific *DNM1L* variants causes differential impact on peroxisomal morphology

To compare the functional effects of disease-associated *DNM1L* variants across protein domains, we generated transgenic *Drosophila* expressing human DRP1 variants representing the GTPase, middle, and GTPase effector domains (GED). These included A192E in the GTPase domain variant associated with autosomal dominant optic atrophy (9), G350R, a middle domain variant we previously reported associated with severe epileptic encephalopathy (16) as well as the L700del in the GED domain. We also included a previously characterized GED variant Y691C (7), allowing comparison of two disease-associated variants within the GED.

We expressed reference human DNM1L (*DNM1L^Ref^)* and lacZ as control or the A192E, G350R, L700del, and Y691C variants in *Drosophila* and examined GFP-SKL-labeled peroxisomes in larval salivary gland cells (**Fig. S3**).

Peroxisomes in cells expressing *DNM1L^Ref^* or the GTPase-domain variant *DNM1L^A192E^* were similar to lacZ controls, appearing as uniformly distributed puncta throughout the cell (**Fig. S3 A–C’**). In contrast, expression of the middle-domain variant *DNM1L^G350R^* produced enlarged and clustered peroxisomes with prominent perinuclear accumulation (**Fig. S3 D, D’**). Expression of *DNM1L^L700del^* resulted in increased variability in peroxisomal size, without an apparent alteration in their intracellular distribution (**Fig. S3 E, E’**). The GED variant *DNM1L^Y691C^* also produced abnormal peroxisomal morphology, including enlarged and clustered peroxisomes and a reduction in peroxisome number (**Fig. S3 F, F’**).

Quantification demonstrated a significant reduction in peroxisome number in flies expressing G350R, L700del, and Y691C, whereas the reduction observed with A192E did not reach statistical significance (**Fig. S3 G**). These findings demonstrate differential effects of disease-associated DNM1L variants on peroxisomal morphology, with greater abnormalities produced by the middle-domain and GED variants than by the GTPase-domain variant A192E.

### *DNM1L* variants rescue the *Drp1* loss of function differentially

To compare the functional activity of the DNM1L variants *in vivo*, we expressed each human transgene ubiquitously using *da-Gal4* in *Drp1¹/Drp1²* mutant flies and assessed rescue of lethality. Expression of *DNM1L^Ref^*has previously been shown to rescue *Drp1* mutant lethality (7) and was used as the reference for comparison. The GTPase-domain variant *DNM1L^A192E^* rescued viability similarly to *DNM1L^Ref^* (**Table 2**). In contrast, the middle-domain variant *DNM1L^G350R^* showed markedly impaired rescue, with approximately 3% viability at 25°C and no surviving adults at 29°C. The GED variants showed intermediate rescue: *DNM1L^Y691C^* produced few surviving adults, whereas *DNM1L^L700del^*resulted in approximately 25% viability at 29°C (**Table 2**).

**Table 2:** Impact on lethality phenotype by overexpressing *DNM1L* variants in Drp1 LOF background.

| **Lethality Rescue of Drp1 null mutant** | ***DNM1L<sup>(REF)</sup>*** | **GD**<br>***DNM1L<sup>(A192E)</sup>*** | **MD**<br>***DNM1L<sup>(G350R)</sup>*** | **GED**<br>***DNM1L<sup>(Y691C)</sup>***<br>***DNM1L<sup>(L700del)</sup>*** |
|---|---|---|---|---|
| **18°C** | Yes | Yes | Few Escapes | Partial |
| **25°C** | Yes | Yes | Few Escapes | Partial |
| **29°C** | Yes | Yes | **Lethal** | Partial |

**Genotype:** *Drp1<sup>1</sup>/Drp1<sup>2</sup> ; da-Gal4/UAS-DNM1L<sup>(X)</sup>*
**X: Human reference/GD (A192E)/ MD (G350R)/ GED (Y691C, L700del)**

We next examined whether these differences in functional rescue were accompanied by differences in peroxisomal morphology in larval muscle (**Fig. 4A-4G**). *DNM1L^Ref^* and A192E-expressing animals exhibited predominantly spherical, uniformly distributed Pex3-positive peroxisomes (**Fig. 4B, B’.& C,C’** respectively). Although peroxisome number was significantly reduced in A192E-expressing animals compared with *DNM1L^Ref^* (**Fig. 4G**), overall peroxisomal morphology was relatively preserved. In contrast, G350R produced severe abnormalities characterized by enlarged, clustered, and tubular peroxisomes (**Fig. 4D, D’)** resembling those observed in *Drp1¹/Drp1²* loss-of-function animals (**Fig. 4A, A’**). The GED variants L700del and Y691C produced intermediate morphological abnormalities, with mixed populations of variably sized and clustered peroxisomes (**Fig. 4E,E’ & F,F’** respectively). Quantitative analysis confirmed significant differences in peroxisome number and size among the variants, with the most severe morphological abnormalities observed with G350R (**Fig. 4G**).

**Fig 4.**
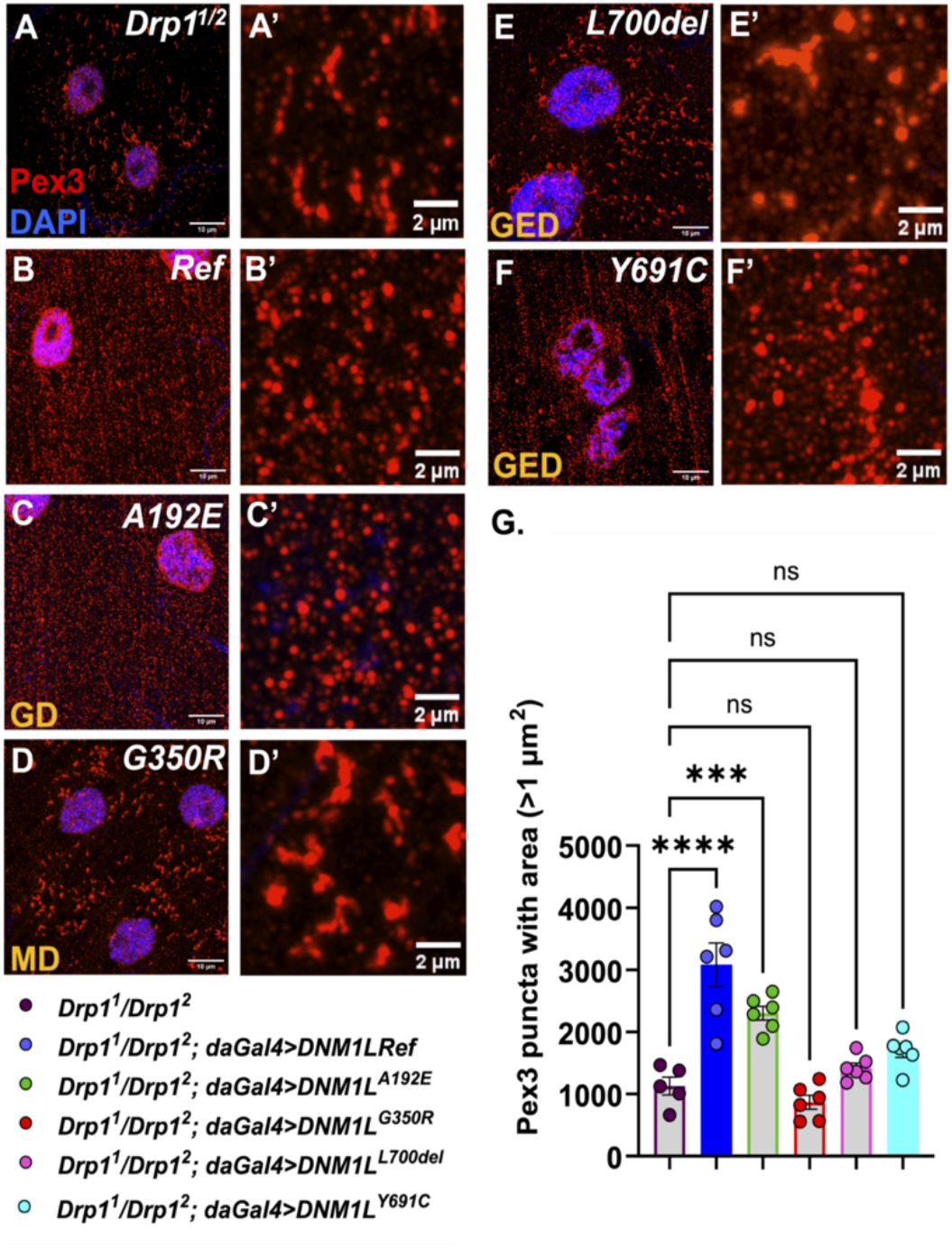
*DNM1L* variants expression in *Drp1* LOF background shows variable degree of rescue and alteration in peroxisomal morphology. **The panel set A, A’ -F, F’** shows representative confocal images illustrating pex3 stained peroxisomes at different magnification (40x Zoom 2.5 & zoom 5 respectively). **A, A’** represents *Drp1^1^/Drp1^2^*, depicting fewer and elongated peroxisomes. B, B’ represents *DNM1L^Ref^*. C, C’ represents GTPase domain (GD) variant A192E that exhibits similar pex3 pattern to the reference group. The middle domain (MD) variant G350R **(D, D’)**, displayed enlarged clustered and tubular peroxisomes with abnormal distribution. GTPase effector domain (GED) variants *L700del* **(E, E’)** and *Y691C* **(F, F’)** displayed enlarged clustered and fewer peroxisomes. **I.** Graphical illustration of Pex3 positive puncta number quantitation. N=5-6 larvae, ***p<0.001;****p<0.0001

Together, the differential rescue of *Drp1* mutant lethality and the severity of peroxisomal abnormalities support a functional allelic series among the *DNM1L* variants examined, with A192E showing minimal functional impairment, G350R the most severe, and the GED variants L700del and Y691C exhibiting intermediate effects.

### *DNM1L* variants produce graded peroxisomal and mitochondrial abnormalities in a *Drp1*-sensitized background

Based on our functional studies of DNM1L variants in the *Drp1¹/Drp1²* loss-of-function background, we next expressed the variants in a heterozygous *Drp1¹/+* sensitized background. Most dominant DNM1L-associated disorders, including the L700del cases described here, result from heterozygous variants acting in the presence of functional DRP1. Although the *Drp1¹/+* model does not precisely reproduce the human genotype, it provides a sensitized genetic background in which endogenous Drp1 function is retained and the relative effects of disease-associated human DNM1L variants can be compared. We therefore used this system to examine the effects of the variants on peroxisomal and mitochondrial morphology in larval muscle.

Using the muscle-specific Mef2-GAL4 driver, we expressed *DNM1L^Ref^*and each disease-associated DNM1L variant in the *Drp1¹/+* background. Peroxisomes in muscles expressing lacZ or *DNM1L^Ref^* appeared predominantly as spherical puncta (**Fig. S4 A, A’ & B, B’** respectively), and similar morphology was observed with the GTPase-domain variant A192E (**Fig. S4 C, C’**). In contrast, expression of the middle-domain variant G350R produced severe peroxisomal abnormalities characterized by elongated and tubular peroxisomes (**Fig. S4 D, D’**). The GED variants L700del and Y691C also altered peroxisomal morphology, but these abnormalities were less severe than those observed with G350R (**Fig. S4 E,E’ & F,F’**). Quantitative analysis demonstrated differences in peroxisome number and morphology among the variants (**Fig. S4 G**). Although A192E reduced peroxisome number, overall peroxisomal morphology was relatively preserved, whereas G350R produced the most severe morphological abnormalities and the GED variants exhibited intermediate phenotypes.

We next examined mitochondrial morphology in the same sensitized background. Muscles expressing lacZ or *DNM1L^Ref^* displayed an elongated, interconnected mitochondrial network (**Fig. S5 A,A’–B,B’)**. A192E produced relatively mild mitochondrial abnormalities (**Fig. S5 C, C’**), whereas G350R caused marked disruption of mitochondrial morphology, with enlarged mitochondria and severe abnormalities of the mitochondrial network (**Fig. S5 D, D’**). L700del and Y691C also produced enlarged mitochondria and altered mitochondrial networks, with phenotypes less severe than those observed with G350R (**Fig. S5 E–F’**). Quantification demonstrated a significant increase in mitochondrial width in flies expressing the middle-domain and GED variants (**Fig. S5 G**), consistent with impaired mitochondrial fission.

Together, analysis of peroxisomal and mitochondrial morphology in the *Drp1*-sensitized background demonstrated graded functional effects among the disease-associated *DNM1L* variants. A192E produced relatively mild abnormalities, G350R produced the most severe defects, and the GED variants L700del and Y691C exhibited intermediate phenotypes. These findings, together with the differential rescue of *Drp1¹/Drp1²* lethality, support a functional allelic spectrum among DNM1L variants affecting different protein domains.

### *DNM1L* variants impair locomotor function and growth in a Drp1-sensitized background

To determine whether the morphological abnormalities associated with disease-associated *DNM1L* variants were accompanied by functional deficits *in vivo*, we assessed larval locomotion following muscle-specific expression of *DNM1L^Ref^*, A192E, G350R, or L700del in the *Drp1¹/+* sensitized background. Larval crawling was quantified by measuring distance traveled, average speed, and maximum speed over one minute (**Fig. 5A**). Expression of the disease-associated variants significantly impaired locomotor performance compared with *DNM1L^Ref^*, including reduced distance traveled (**Fig. 5B**), average speed (**Fig. 5C**), and maximum speed (**Fig. 5D**).

**Fig 5.**
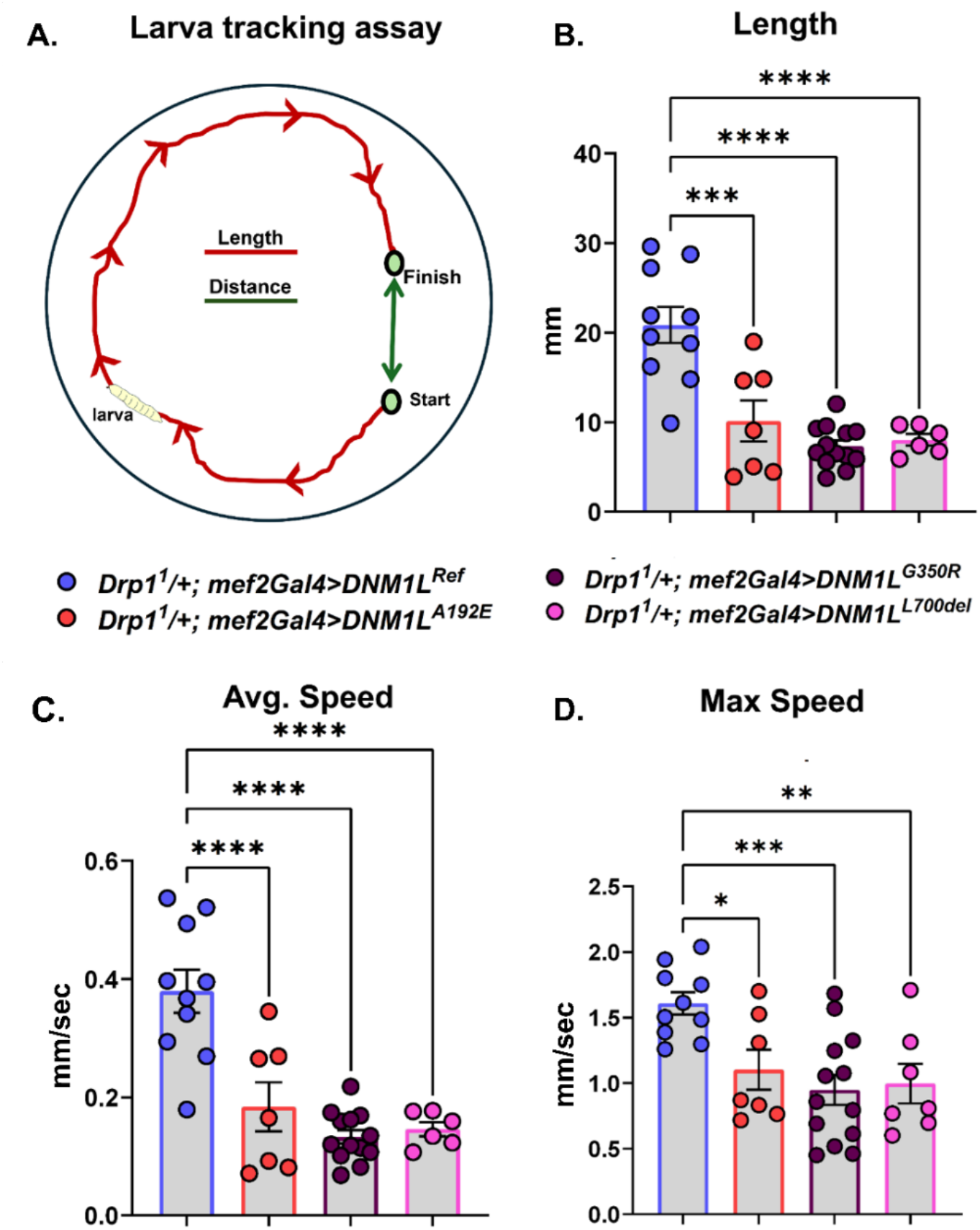
Overexpressing *DNM1L* variants in muscle with Drp1 sensitized background display significant locomotion defect in variant larva. A. illustrates the larva tracking assay done in the plate containing grape juice agar media. B: The graph shows the length travelled during a given time, variants length coverage was significantly down as compared to Ref. C: illustrates the average speed of the larva, significantly down among variants. D: Graph illustrates the max speed acquire by the larva, variants display significantly lower max speed, GD variant A192E showed milder effect, while MD variant G350R showed strong impact and GED variant L700del showed intermediate impact on max speed.

We also assessed pupal length as a measure of organismal growth. Flies expressing the disease-associated DNM1L variants exhibited significantly reduced pupal length compared with controls (**Fig. S6**). Thus, expression of disease-associated DNM1L variants in muscle produces measurable organismal phenotypes, including impaired locomotor performance and reduced growth.

### *DNM1L* variants produce graded abnormalities in peroxisomal morphology in glial cells

Because *DNM1L*-associated disorders predominantly affect the nervous system, we next examined the effects of the disease-associated variants on peroxisomal morphology in glial cells. Each human DNM1L transgene was expressed using the glial-specific repo-GAL4 driver in the *Drp1¹/+* sensitized background, and peroxisomal morphology was examined in the larval brain (**Fig. 6**).

**Fig 6.**
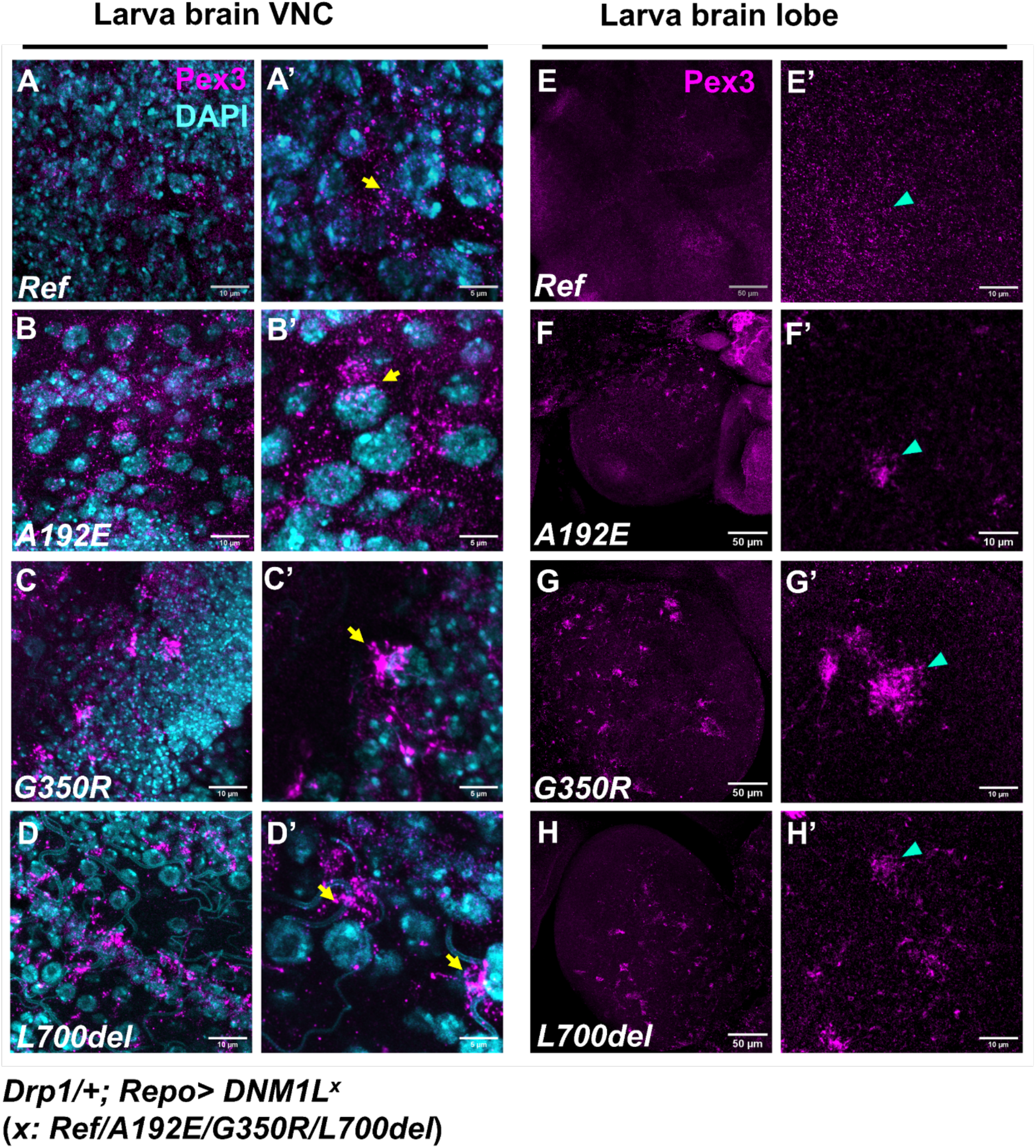
Overexpressing *DNM1L* variants with Drp1 sensitized background in glial cells exhibits clustering of peroxisomes in larva brain. Representative images depicting peroxisomal phenotype in larva brain VNC (A, A’ – D, D’) and larva brain lobes (BL) (E, E’ – H, H’). The Ref group exhibits uniform distribution of peroxisomes immuno-stained with pex3 in both VNC (A, A’) and BL (E, E’) region of the larva brain. The GD variant A192E exhibits near normal distribution of peroxisomes with mild clustering both in VNC (B, B’) and BL (F, F’). The MD variant G350R exhibits branches like clustering uniquely on one side of nuclei in VNC (C, C’), while multiple clustered peroxisomes patches were observed throughout the BL (G, G’). The GED variants *L700del* exhibit somewhat similar but milder clustering of peroxisomes at one side of the nuclei in the VNC (D, D’), while milder peroxisomes cluster patches were observed in BL (H, H’).

Glial cells expressing *DNM1L^Ref^* exhibited predominantly spherical peroxisomes that were uniformly distributed throughout the ventral nerve cord and brain lobes (**Fig. 6A, A’, E, E’**). Expression of the GTPase-domain variant A192E produced relatively mild peroxisomal clustering in the ventral nerve cord (**Fig. 6B, B’**). In contrast, the middle-domain variant G350R produced prominent clusters of elongated, tubular peroxisomes localized near the nuclear periphery (**Fig. 6C, C’**). The GED variant L700del produced a similar but less pronounced clustering phenotype (**Fig. 6D, D’**). Examination of the brain lobes similarly demonstrated increasing peroxisomal abnormalities across the GTPase-domain, GED, and middle-domain variants (**Fig. 6**).

Thus, the three representative DNM1L variants produced graded effects on peroxisomal morphology in glial cells, with A192E producing relatively mild abnormalities, L700del an intermediate phenotype, and G350R the most severe phenotype. The persistence of this functional hierarchy in glia further supports a relationship between *DNM1L* functional domain, allelic severity, and organelle dysfunction.

## Discussion

Autosomal dominant pathogenic variants in *DNM1L* are associated with a broad spectrum of neurological disorders, ranging from isolated optic atrophy to severe developmental and epileptic encephalopathy (9, 22). Emerging genotype–phenotype data suggest that variants affecting different functional domains of DRP1 may differ in their clinical and functional severity, although variants affecting the GTPase effector domain (GED) remain comparatively poorly characterized. Here, we describe four individuals harboring a recurrent *L700del* GED variant who share a neurological phenotype with early-onset dystonia, tremor, gait abnormalities, and motor delay. In contrast to the severe infantile encephalopathy frequently associated with pathogenic middle-domain variants, none of these individuals experienced infantile seizures or severe infantile encephalopathy. These findings expand the recognized phenotypic spectrum associated with *DNM1L* variants and further implicate the GED in a movement-disorder-predominant disease.

Functional studies in patient-derived fibroblasts support the pathogenicity of *L700del* and demonstrate abnormalities of both DRP1-regulated organelles. *L700del* fibroblasts exhibited elongated peroxisomes, reduced peroxisome number, and abnormal mitochondrial morphology. Fibroblasts from two affected individuals also showed impaired growth under conditions that increase dependence on mitochondrial oxidative metabolism. These findings establish cellular consequences of the recurrent L700del variant and provide a basis for comparing its functional severity with *DNM1L* variants associated with other neurological phenotypes.

Our studies in *Drosophila* allowed disease-associated *DNM1L* variants to be directly compared within a common genetic background across multiple functional assays. Across both the *Drp1¹/Drp1²* loss-of-function and *Drp1¹/+* sensitized backgrounds, the variants produced graded effects on viability and mitochondrial and peroxisomal morphology. The sensitized *Drp1¹/+* background is particularly informative because it permits variant effects to be assessed in the presence of residual endogenous Drp1 function, providing an experimental approximation of the heterozygous state associated with dominant human *DNM1L* disease. The GTPase-domain variant A192E generally produced the mildest abnormalities, whereas the middle-domain variant G350R produced the most severe defects. L700del consistently exhibited intermediate functional effects, in parallel with the intermediate clinical severity observed in the individuals described here. Interestingly, comparison of L700del with the previously reported GED variant Y691C (7), demonstrated that variants within the same domain do not necessarily have equivalent functional effects. Y691C exhibited substantial variability across assays, including pronounced effects on peroxisomal morphology and viability, consistent with the more severe neurological phenotype previously reported for this variant, which included epilepsy and absent verbal communication. Thus, our findings support a functional allelic spectrum in *DNM1L*-associated disease in which protein domain contributes to functional severity, yet variant-specific effects within individual domains further modify the resulting phenotype.

The functional differences observed among disease-associated *DNM1L* variants are consistent with the distinct roles of DRP1 domains in the fission cycle. The GTPase domain mediates GTP binding and hydrolysis, whereas the middle domain contributes to higher-order assembly of DRP1 complexes required for membrane constriction (1, 23). The GED participates in regulation of DRP1 assembly and GTPase activity, and pathogenic GED variants can disrupt these functions without completely abolishing DRP1 activity (24). Thus, variants affecting different domains may alter distinct steps in DRP1-mediated fission and produce different degrees of functional impairment (24). Our direct comparison of disease-associated variants extends these structure-function observations by demonstrating that differences in allelic strength can be detected across multiple *in vivo* assays and broadly parallel differences in human neurological severity.

Allelic strength alone may not fully explain the distinct neurological phenotypes associated with DNM1L variants, as the consequences of impaired organelle dynamics are also likely to vary among cell types and tissues. In our Drosophila model, disease-associated DNM1L variants produced graded abnormalities of peroxisomal morphology in glial cells, with relatively mild effects of A192E, intermediate effects of L700del, and severe abnormalities with G350R. These findings raise the possibility that tissue-specific thresholds for mitochondrial and peroxisomal dysfunction contribute to the phenotypic diversity of DNM1L-associated disease. Glial cells have important roles in neuronal metabolic support and in the response to oxidative and lipid stress (25, 26), suggesting that disruption of mitochondrial and peroxisomal dynamics in glia could contribute to neurological disease. In addition, DRP1 recruitment and activation involve overlapping but distinct organelle-specific machinery, including MFF, MiD49, and MiD51 at mitochondria and PEX11β and MFF at peroxisomes (2). Differences in these regulatory networks among organelles and cell types could further modify the consequences of individual DNM1L variants.

Finally, patient fibroblasts harboring the DNM1L L700del variant exhibited a selective growth defect in galactose-containing medium, while growth remained comparable to control fibroblasts in glucose-containing medium. Because galactose limits glycolytic ATP production and increases cellular dependence on mitochondrial oxidative metabolism, this finding reveals a metabolic vulnerability associated with the L700del variant. Importantly, bezafibrate partially rescued this growth defect. This finding is consistent with previous studies demonstrating beneficial effects of bezafibrate in DNM1L-mutant fibroblasts (21) and suggests that at least some of the metabolic consequences of DNM1L dysfunction are pharmacologically modifiable. While this is not evidence for a near-term therapy, this data provides a proof-of-principle demonstration that the cellular dysfunction can be modified. Although additional studies will be required to define the mechanism of rescue and determine whether these findings translate beyond cultured fibroblasts, our results provide proof-of-concept for investigating metabolic modulation as a potential therapeutic approach in DNM1L-associated disease.

In conclusion, the recurrent DNM1L L700del variant is associated with a distinctive movement-disorder phenotype characterized by early motor delay, dystonia, tremor, and gait abnormalities without the severe infantile encephalopathy characteristic of many middle-domain DNM1L variants. Functional studies in patient-derived cells and Drosophila demonstrate that L700del disrupts both mitochondrial and peroxisomal dynamics and exhibits intermediate functional severity within a broader spectrum of disease-associated DNM1L alleles. These findings support a model in which differences in allelic strength, together with tissue-specific vulnerability to impaired organelle dynamics, contribute to the striking clinical heterogeneity of DNM1L-associated disease. The metabolic vulnerability identified in L700del fibroblasts and its partial rescue by bezafibrate further suggest that downstream consequences of DNM1L dysfunction may be pharmacologically modifiable. Together, these findings provide a framework connecting DNM1L genotype, functional severity, organelle dysfunction, and clinical phenotype.

## Acknowledgments

The authors sincerely thank the family for their participation in this study. We are also grateful to the *DNM1L* Foundation for their support. Special thanks to Professor James McNew (Rice University) for generously providing the Pex3 antibody used in this work.

## Supplementary figures

**Figure S1:**
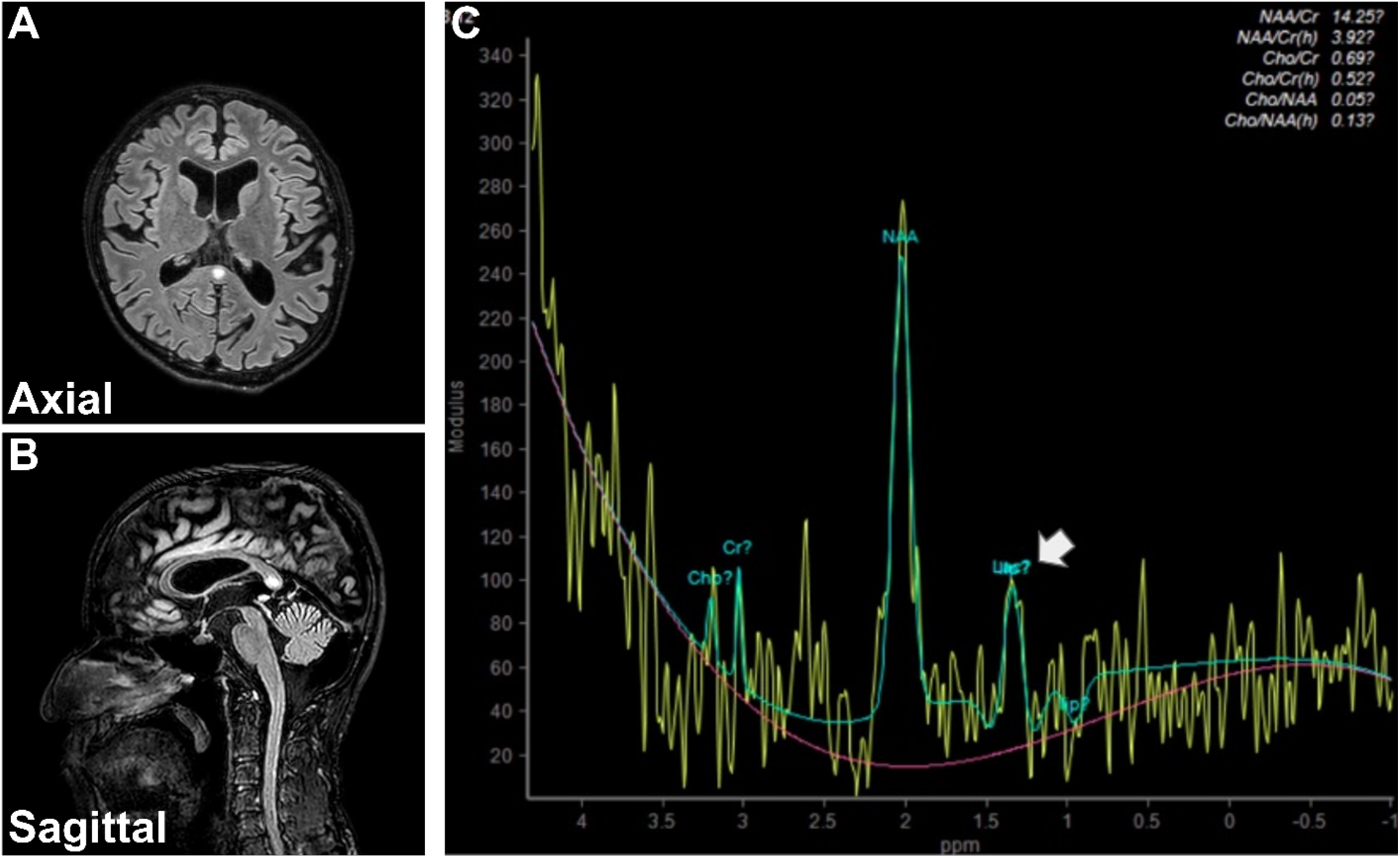
Brain MRI (T2-FLAIR) demonstrates a 0.7-cm splenial lesion with diffusion restriction and no enhancement (CLOCC-like), associated with moderate brain volume loss and a prominent retrocerebellar CSF space. Intralesional single-voxel long-TE spectroscopy shows a lactate peak (white arrow) with otherwise normal metabolite ratios.

**Fig S2:**
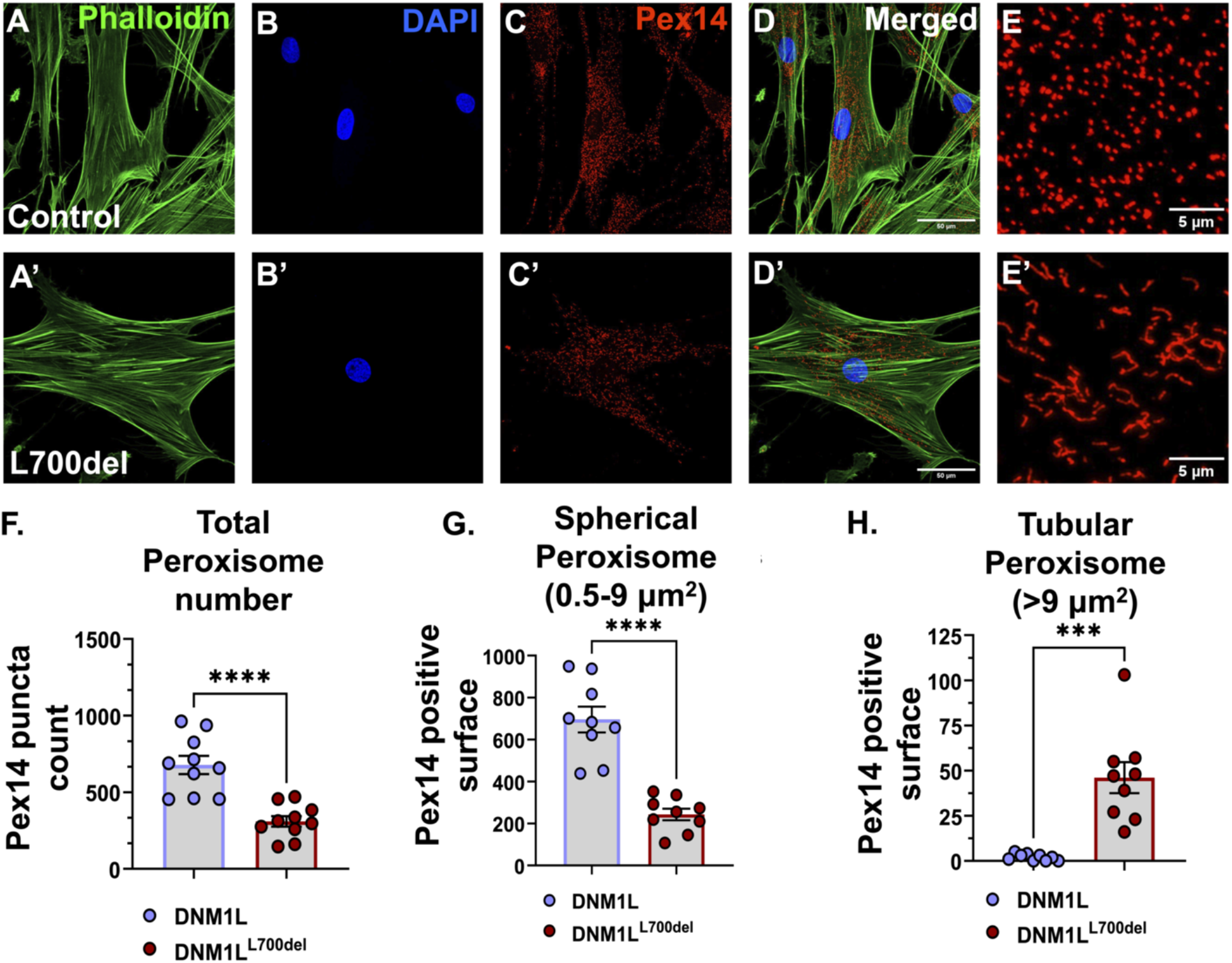
Impact of *DNM1L^L700del^*variant on peroxisomal morphology. Illustration of peroxisomal morphology based on Pex14 immunostaining in control versus patient skin fibroblast cells (A & A’ respectively). The fibroblast from control (A-D, E) exhibits spherical shaped peroxisomes (A, E), while patient fibroblast with *DNM1L^L700del^* variant exhibited tubular and elongated peroxisomes (A’, E’). The total peroxisome number, and number of tubulated and non-tubulated peroxisomes were quantified using IMARIS (F-H). Total peroxisome number was decreased in *DNM1L^L700del^*variant contrary to control (F). Number of non-tubulated peroxisomes was also down in fibroblast from *DNM1L^L700del^* variant versus control (G). The number of tubulated peroxisomes was significantly high in fibroblast from *DNM1L^L700del^* variant as compared to control (H). The tubular elongated peroxisomes depict failure in fission process. Student t-test was performed for statistical significance ***p=0.0002, ****p<0.0001

**Figure S3.**
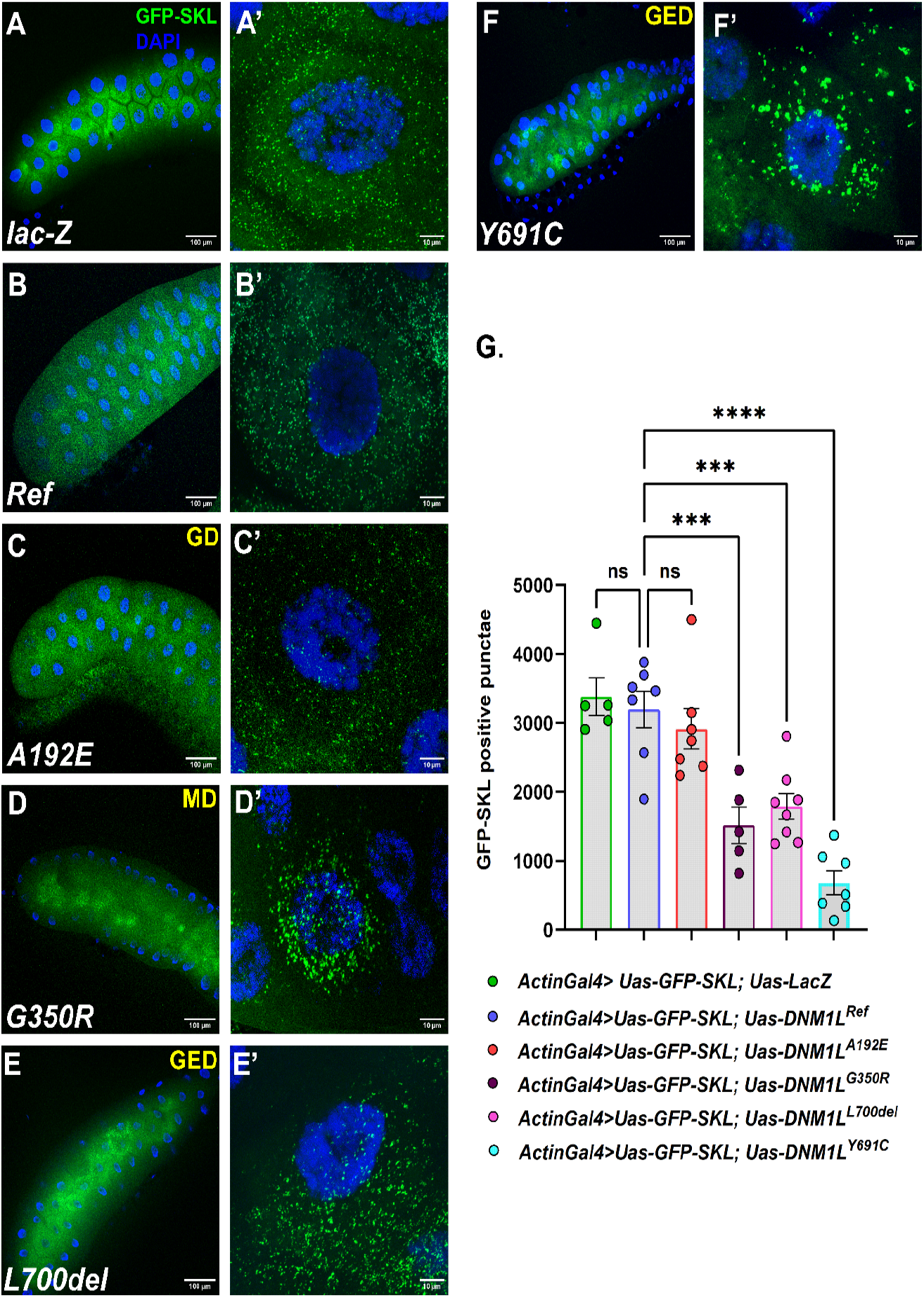
Overexpression of *DNM1L* variant, with GFP-SKL in salivary gland affects peroxisome number and morphology. **Panel A-F** represents confocal images of larval salivary glands (SG) captured using a 20x objective with scan zoom 0.6 (scale bar = 100 µm), and **Panel A’-F’** represents confocal images of single corpuscular cell of SG acquired using 63x water objective with scan zoom 1.5 (scale bar = 10 µm). The Peroxisome tag GFP-SKL is shown in green while nucleus was stained with DAPI (blue). **Panel A-A’** represents the control (*lacZ* overexpression), while **B-B’** shows overexpression of the human reference *DNM1L^Ref^*. **Panels C, C’-F, F’,** correspond to domain-specific *DNM1L* variants. The observed phenotypes vary among groups. The GTPase domain (GD) variant A192E **(C’)** displayed no remarkable effect on peroxisomal size and appeared like the control. The most severe phenotype was observed with middle domain (MD) variants G350R **(D’)**, which displayed enlarged peroxisomes with abnormal distribution. GTPase effector domain (GED) variants *L700del* **(E’)**, showed moderate peroxisomal defect, while *Y691C* **(F’)** displayed enlarged peroxisomes with abnormal distribution while. and human reference *DNM1L* groups. **H** Graphical illustration of peroxisomal puncta number derived from GFK-SKL puncta analysis through surface module of IMARIS v9.8. N=5-7 animals from each group. ***p=0.0002, ****p<0.0001

**Fig S4.**
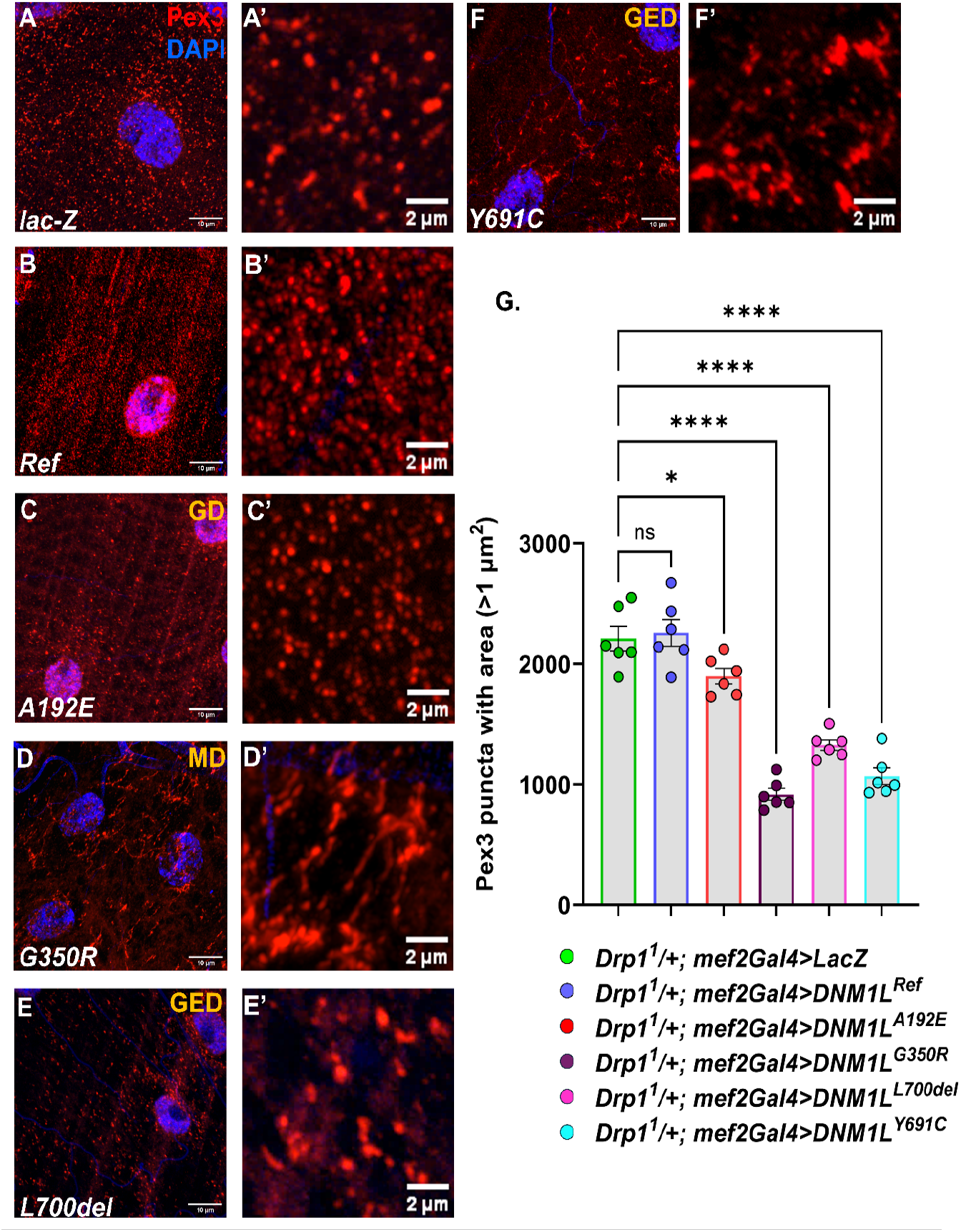
Overexpression of *DNM1L* variants in *Drp1*-sensitized larval muscle affects peroxisomal morphology and number to varying degrees. Representative confocal images of larval body wall muscle stained with Pex3 (peroxisomes, red) and DAPI (nuclei, blue) are shown. **Panels A-F** were captured using a 40x objective with scan zoom 2.5 (scale bar = 10 µm), and **panels A’-F’** were acquired with scan zoom 5 (scale bar = 2 µm). **Panel A-A’** represents control (*lacZ* overexpression), while **B-B’** shows overexpression of the *DNM1L^Ref^*. **Panels C, C’-F-F’** correspond to domain-specific *DNM1L* variants. The observed phenotypes vary in severity and are categorized as severe, mild, or negligible. The GTPase domain (GD) variant A192E **(C, C’)** displayed peroxisomal morphology like the control and human reference *DNM1L* groups. The most severe phenotype was observed with middle domain (MD) variants G350R **(D, D’)**, which display pronounced peroxisomal tubulation and clustering associated morphological abnormalities. Less severe, or mild, phenotypes were observed with GTPase effector domain (GED) variants *L700del* **(E, E’)** and *Y691C* **(F, F’)**, which showed moderate peroxisomal defects. **G.** Quantification of peroxisomal number, represented as Pex3-positive puncta count, revealed that all *DNM1L* variants caused a significant reduction in peroxisomal number compared to the control group. Statistical analysis was performed with **N = 6** for each group,**\*\*\*\*p < 0.0001, *p < 0.05.**

**Fig S5.**
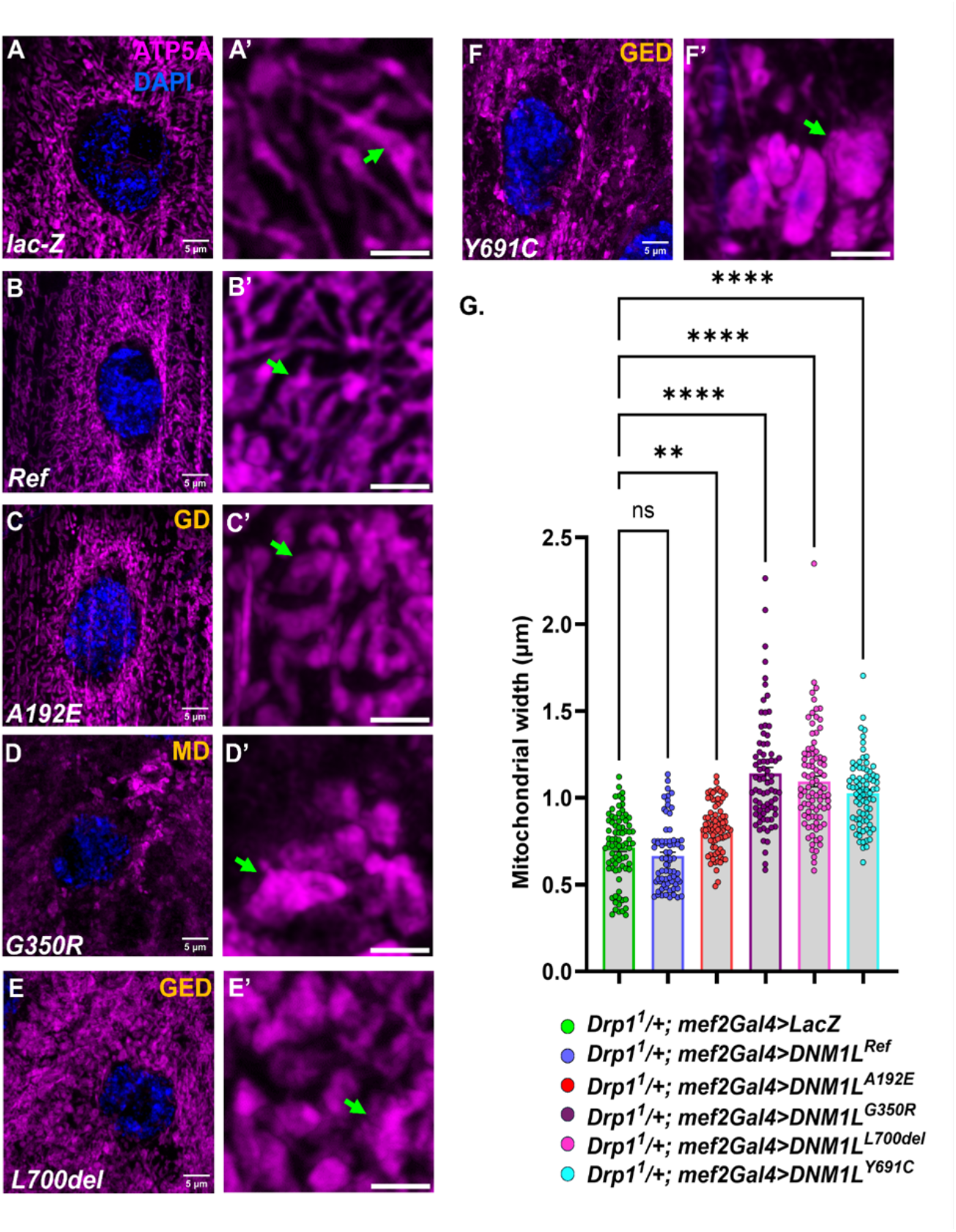
Overexpression of *DNM1L* variants in *Drp1*-sensitized larval muscle alters mitochondrial morphology from tubular to more hyperfused & fragmented. Representative confocal images of larval body wall muscle stained with ATP5A (mitochondria complex-V, magenta) and DAPI (nuclei, blue) are shown. **Panels A-F** were captured using a 63x objective with scan zoom 2.5 (scale bar = 5 µm) and **panels A’-F’** were acquired with scan zoom 5 (scale bar = 2 µm). **Panel A-A’** represents control (*lacZ* overexpression), while **B-B’** shows overexpression of the *DNM1L^Ref^*. **Panels C-F, C’-F’** correspond to domain-specific *DNM1L* variants. Mitochondria appear to exhibit more uniform distribution with connected tubular morphology in control **(A, A’)** and *DNM1L^Ref^* group **(B, B’)**, GD variant *A192E* appears to exhibit less connectivity and mitochondria appears less tubular and more swollen **(C, C’)**. The MD variant *G350R* **(D, D’)** displayed pronounced mitochondrial swelling (hyperfused), tubular network loss and morphological abnormalities. Similar but milder phenotypes were observed with GED variants *L700del* **(E, E’)** and *Y691C* **(F, F’)**. **G** displays graphical illustration of mitochondrial width quantitation. N=50+ mitochondria from each group. **p=0.007; ****p<0.0001

**Figure S6.**
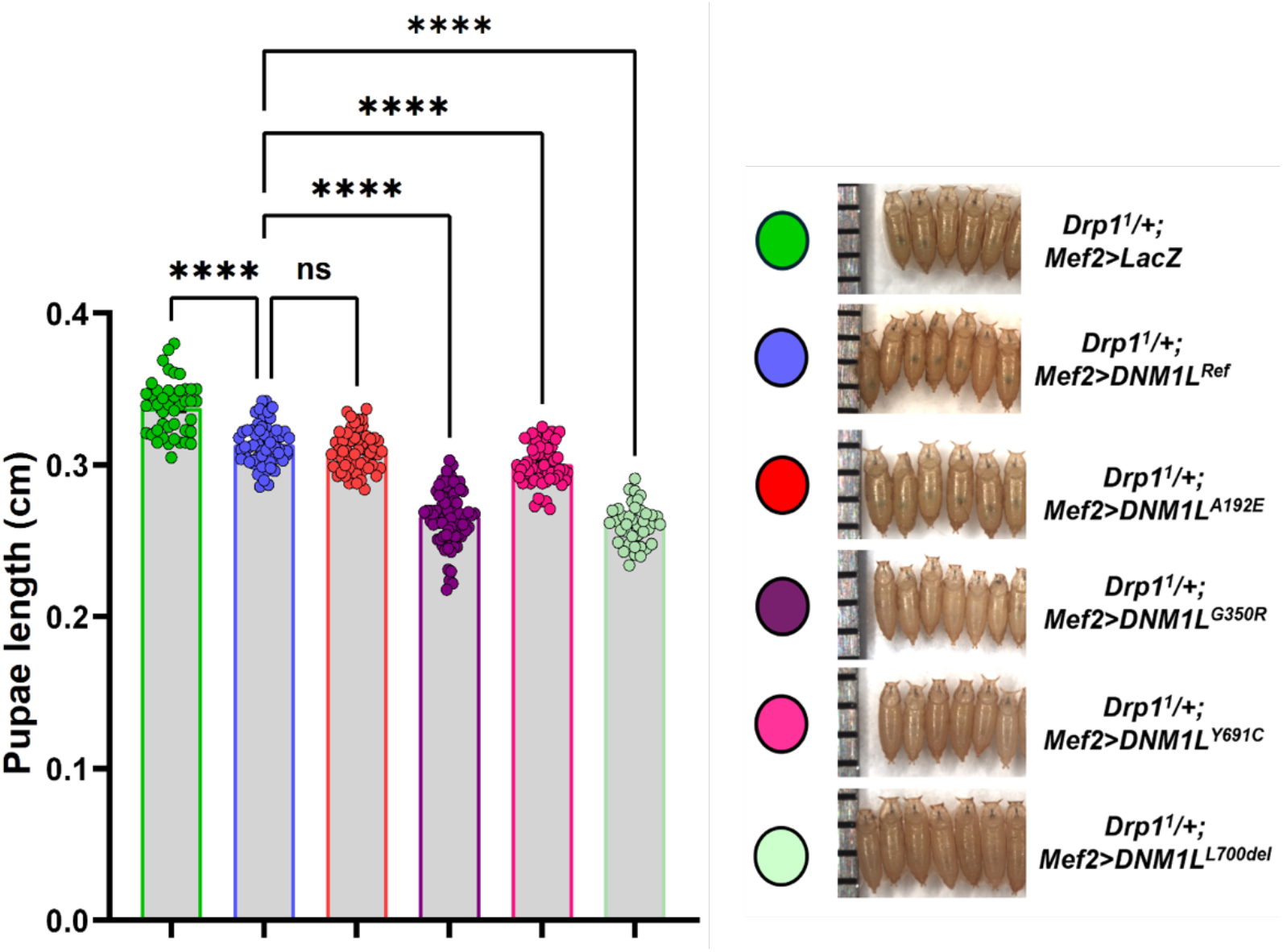
Pupal length in Drp1 sensitized larvae expressing DNM1L ref/variants in larva body wall muscle using Mef2-Gal4 driver.

## References

1. Chang C-R, Blackstone C. Dynamic regulation of mitochondrial fission through modification of the dynamin-related protein Drp1. Ann N Y Acad Sci. 2010;1201(1):34–39.

2. Schrader M, Bonekamp NA, Islinger M. Fission and proliferation of peroxisomes. Biochim Biophys Acta. 2012;1822(9):1343–1357.

3. Koirala S, et al. Molecular architecture of a dynamin adaptor: implications for assembly of mitochondrial fission complexes. J Cell Biol. 2010;191(6):1127–1139.

4. Tilokani L, et al. Mitochondrial dynamics: overview of molecular mechanisms. Essays Biochem. 2018;62(3):341–360.

5. Kamerkar SC, et al. Dynamin-related protein 1 has membrane constricting and severing abilities sufficient for mitochondrial and peroxisomal fission. Nat Commun. 2018;9(1):5239.

6. Imoto Y, Itoh K, Fujiki Y. Molecular basis of mitochondrial and peroxisomal division machineries. Int J Mol Sci. 2020;21(15):E5452.

7. Assia Batzir N, et al. De novo missense variant in the GTPase effector domain (GED) of DNM1L leads to static encephalopathy and seizures. Cold Spring Harb Mol Case Stud. 2019;5(3). 10.1101/mcs.a003673.

8. Fröhlich C, et al. Structural insights into oligomerization and mitochondrial remodelling of dynamin 1-like protein. EMBO J. 2013;32(9):1280–1292.

9. Gerber S, et al. Mutations in DNM1L, as in OPA1, result in dominant optic atrophy despite opposite effects on mitochondrial fusion and fission. Brain. 2017;140(10):2586–2596.

10. Zhang Z, et al. A novel variant of DNM1L expanding the clinical phenotypic spectrum: a case report and literature review. BMC Pediatr. 2024;24(1):104.

11. Yoon G, et al. Lethal Disorder of Mitochondrial Fission Caused by Mutations in DNM1L. J Pediatr. 2016;171:313–6.e1–2.

12. Nasca A, et al. Biallelic Mutations in DNM1L are Associated with a Slowly Progressive Infantile Encephalopathy. Hum Mutat. 2016;37(9):898–903.

13. Vanstone JR, et al. DNM1L-related mitochondrial fission defect presenting as refractory epilepsy. Eur J Hum Genet. 2016;24(7):1084–1088.

14. Sheffer R, et al. Postnatal microcephaly and pain insensitivity due to a de novo heterozygous DNM1L mutation causing impaired mitochondrial fission and function. Am J Med Genet A. 2016;170(6):1603–1607.

15. Verstreken P, et al. Synaptic mitochondria are critical for mobilization of reserve pool vesicles at Drosophila neuromuscular junctions. Neuron. 2005;47(3):365–378.

16. Chao Y-H, et al. Missense variants in the middle domain of DNM1L in cases of infantile encephalopathy alter peroxisomes and mitochondria when assayed in Drosophila. Hum Mol Genet. 2016;25(9):1846–1856.

17. Venken KJT, et al. P[acman]: a BAC transgenic platform for targeted insertion of large DNA fragments in D. melanogaster. Science. 2006;314(5806):1747–1751.

18. Wangler MF, et al. Peroxisomal biogenesis is genetically and biochemically linked to carbohydrate metabolism in Drosophila and mouse. PLoS Genet. 2017;13(6):e1006825.

19. Faust JE, et al. Peroxisomes are required for lipid metabolism and muscle function in Drosophila melanogaster. PLoS One. 2014;9(6):e100213.

20. Brooks DS, et al. Optimization of wrMTrck to monitor Drosophila larval locomotor activity. J Insect Physiol. 2016;93–94:11–17.

21. Douiev L, et al. Bezafibrate improves mitochondrial fission and function in DNM1L-deficient patient cells. Cells. 2020;9(2):E301.

22. Fahrner JA, et al. A novel de novo dominant negative mutation in DNM1L impairs mitochondrial fission and presents as childhood epileptic encephalopathy. Am J Med Genet A. 2016;170(8):2002–2011.

23. Rochon, Kristy, et al. “Structural basis for regulated assembly of the mitochondrial fission GTPase Drp1.” Nature communications 15.1 (2024): 1328

24. Nolden KA, et al. Novel DNM1L variants impair mitochondrial dynamics through divergent mechanisms. Life Sci Alliance. 2022;5(12). 10.26508/lsa.202101284.

25. Lee KH, Cha M, Lee BH. Crosstalk between neuron and glial cells in oxidative injury and neuroprotection [preprint]. Int J Mol Sci. 2021;22(24). 10.3390/ijms222413315.

26. Barber CN, Raben DM. Lipid metabolism crosstalk in the brain: Glia and neurons [preprint]. Front Cell Neurosci. 2019;13. 10.3389/fncel.2019.00212.

